# Actin-dependent recruitment of AGO2 to the zonula adherens

**DOI:** 10.1101/2022.03.10.483874

**Authors:** Mary Catherine Bridges, Joyce Nair-Menon, Alyssa Risner, Douglas W. Jimenez, Amanda C. Daulagala, Christina Kingsley, Madison E. Davis, Antonis Kourtidis

**Affiliations:** Department of Regenerative Medicine and Cell Biology, Medical University of South Carolina, 173 Ashley Avenue, Charleston, SC 29425, USA

## Abstract

Adherens junctions are cadherin-based structures critical for cellular architecture. E-cadherin junctions in mature epithelial cell monolayers tether to an apical actomyosin ring to form the zonula adherens (ZA). We have previously shown that the adherens junction protein PLEKHA7 associates with and regulates the function of the core RNA interference (RNAi) component AGO2 specifically at the ZA. However, the mechanism mediating Ago2 recruitment to the ZA remained unexplored. Here, we reveal that this ZA-specific recruitment of AGO2 depends on both the structural and tensile integrity of the actomyosin cytoskeleton. We found that depletion of not only PLEKHA7, but also either of three PLEKHA7-interacting, LIM-domain family proteins, namely LMO7, LIMCH1, and PDLIM1, results in disruption of actomyosin organization and tension, as well as disruption of AGO2 junctional localization and of its miRNA-binding ability. We also show that AGO2 binds Myosin IIB and that PLEKHA7, LMO7, LIMCH1, and PDLIM1 all disrupt interaction of AGO2 with Myosin IIB at the ZA. These results demonstrate that recruitment of Ago2 to the ZA is sensitive to actomyosin perturbations, introducing the concept of a mechanosensitive RNAi machinery, with potential implications in tissue remodeling and in disease.

**Summary:** Recruitment and miRNA-binding activity of the key RNA interference (RNAi) component AGO2 to epithelial zonula adherens depends on apical actomyosin integrity and tension, revealing the existence of a mechanosensitive RNAi machinery at the zonula adherens.

**Significance Statement:**

- Previous work has shown that PLEKHA7 recruits core RNAi components, including AGO2, to regulate tumor-suppressing miRNAs specifically at the zonula adherens (ZA), through an unknown mechanism.
- Here, the authors show that three LIM domain-containing proteins, LMO7, LIMCH1, and PDLIM1, are also responsible for AGO2’s recruitment and miRNA activity at the ZA and that all four PLEKHA7, LMO7, LIMCH1, PDLIM1 mediate AGO2 recruitment to the ZA not due to their protein-protein interactions, but through stabilizing actomyosin structure and tension.
- These findings introduce a mechanosensitive RNAi machinery responsive to actomyosin perturbations, with potentially broad implications in regulation of cellular plasticity.

## INTRODUCTION

The Adherens Junction (AJ) is a cell-cell adhesive structure essential for epithelial tissue organization and homeostasis (Harris and Tepass, 2010; Meng and Takeichi, 2009; Takeichi, 2014). E-cadherin is the core AJ component that forms extracellular linkages between neighboring cells through calcium-dependent homotypic binding (Takeichi, 1995). Intracellularly, E-cadherin binds to β-catenin and p120 catenin (p120) (Reynolds et al., 1994), with p120 playing a key role in stabilizing cadherin junctions (Thoreson et al., 2000). E-cadherin-based junctions also connect to the actin cytoskeleton through α-catenin (Mege and Ishiyama, 2017), a β-catenin binding partner. In polarized epithelia, these connections mature at an apically localized structure known as the zonula adherens (ZA), which links to a circumferential actin ring (Farquhar and Palade, 1963; Meng and Takeichi, 2009). At the ZA, actin organization is intimately involved in the dynamic interplay between contractile force sensation and junctional stabilization (Kovacs et al., 2002; Kovacs et al., 2011; Leerberg et al., 2014; Priya et al., 2013; Ratheesh and Yap, 2012; Shewan et al., 2005; Smutny et al., 2010; Verma et al., 2012; Yamada and Nelson, 2007).

In addition to their critical architectural role, epithelial AJs mediate numerous signaling pathways directing cell behavior (Daulagala et al., 2019; Kourtidis et al., 2017a; Kourtidis et al., 2013; Mendonsa et al., 2018; Ramirez Moreno and Bulgakova, 2021; Salvi and DeMali, 2018; Yu and Elble, 2016; Yulis et al., 2018). This aspect of AJ biology implicates loss of AJ integrity to the progression of multiple diseases, such as cancer (Daulagala et al., 2019; Kourtidis et al., 2017a; Kourtidis et al., 2013; Mendonsa et al., 2018; Ramirez Moreno and Bulgakova, 2021; Yu and Elble, 2016). Along these lines, we have previously demonstrated that cadherin junctions recruit core complexes of RNA interference (RNAi) machinery, such as the microprocessor, the DICER complex, and the RNA-induced Silencing Complex (RISC), to locally regulate microRNA (miRNA) processing and function (Kourtidis et al., 2017b; Kourtidis et al., 2015). Loading of miRNAs onto Argonaute 2 (AGO2), the main enzymatic component of the RISC, leads to translational repression or degradation of target mRNAs (Hammond et al., 2001; Huntzinger and Izaurralde, 2011; Liu et al., 2004; Meister et al., 2004). While AGO2 has been shown to be primarily cytoplasmic, its recruitment to distinct subcellular loci is a mechanism by which RISC activity can be regulated (Antoniou et al., 2014; Bose et al., 2020; Bridge et al., 2017; Detzer et al., 2011; Gagnon et al., 2014; James et al., 2010; Leung et al., 2006; Zeng et al., 2008). Our previous work identified recruitment of AGO2 specifically to the ZA of well-differentiated epithelial cells and tissues, but not at basolateral areas of cell-cell contact (Kourtidis et al., 2017b; Nair-Menon et al., 2020). This recruitment was mediated by PLEKHA7 (Kourtidis et al., 2017b; Kourtidis et al., 2015; Nair-Menon et al., 2020), a ZA-specific p120-binding partner (Meng et al., 2008; Pulimeno et al., 2010). We have shown that loss of junctional AGO2 localization negatively impacts AGO2-loading of a specific set of miRNAs and several of their target oncogenic mRNAs, consequently releasing RISC-mediated oncogene suppression (Kourtidis et al., 2017b; Kourtidis et al., 2015). Furthermore, we demonstrated that junctional localization of AGO2 is broadly lost in human colon tumors and in poorly differentiated colon cancer cell lines, even when cadherin junctions were still present, although not mature, implying for the existence of a mechanism fine-tuning recruitment of AGO2 to mature ZA of well-differentiated epithelia (Nair-Menon et al., 2020).

Indeed, while PLEKHA7 is required for junctional localization of AGO2, what determines AGO2’s specific recruitment to the ZA remained unclear. PLEKHA7 was originally characterized for its role in tethering the minus ends of the microtubules to the ZA (Meng et al., 2008). Notably, a significant body of work demonstrates that intracellular localization of RNA-binding complexes is mediated by the microtubule cytoskeleton (Denes et al., 2021; Kanai et al., 2004; Liao et al., 2019; Pichon et al., 2021; Scarborough et al., 2021). These observations led us to originally hypothesize that AGO2 recruitment to the ZA could be microtubule-directed. Surprisingly, our previous experimentation showed that AGO2, or even PLEKHA7 localization at the ZA is microtubule-independent (Kourtidis et al., 2017b), leaving the mechanism of AGO2 recruitment to the ZA an open question. To obtain clues about this this mechanism, we re-examined in this study our published PLEKHA7 proteome. This re-examination led us to identify three LIM domain-containing proteins as also being responsible for the recruitment of AGO2 to the ZA, in addition to PLEKHA7, and to discover that all four proteins mediate this junctional recruitment of AGO2 through stabilizing the actomyosin cytoskeleton at the ZA.

## RESULTS

### PLEKHA7 and AGO2 associate with LIM domain-containing proteins at the ZA

To obtain insight on the potential mechanisms involved in AGO2 recruitment to the ZA, we re-examined our published dataset of PLEKHA7-interacting proteins (publicly available at http://www.imexconsortium.org/; identifier: IM-25739) (Kourtidis et al., 2017b). In that experiment, PLEKHA7 immunoprecipitation from Caco2 well-differentiated intestinal epithelial cells and subsequent mass-spectrometry analysis identified more than 600 potential PLEKHA7 interactors (Kourtidis et al., 2017b). We have reported that a significant set of these interactors are RNA-binding proteins, such as RISC components (Kourtidis et al., 2017b). Here, we performed gene ontology (GO) enrichment analysis on this dataset. This analysis revealed that in addition to RNA-binding proteins, the set of PLEKHA7-associated proteins is significantly enriched for processes and functions related to the actin cytoskeleton (Figure 1, A and B). This result aligns with our previous findings that PLEKHA7 stabilizes apical actin and that it associates with certain actin-binding proteins (Kourtidis and Anastasiadis, 2016; Kourtidis et al., 2015). Overall, the GO analyses point to RNA-binding and actin cytoskeletal-mediated functions as the top roles in which PLEKHA7 is potentially involved.

**FIGURE 1.**
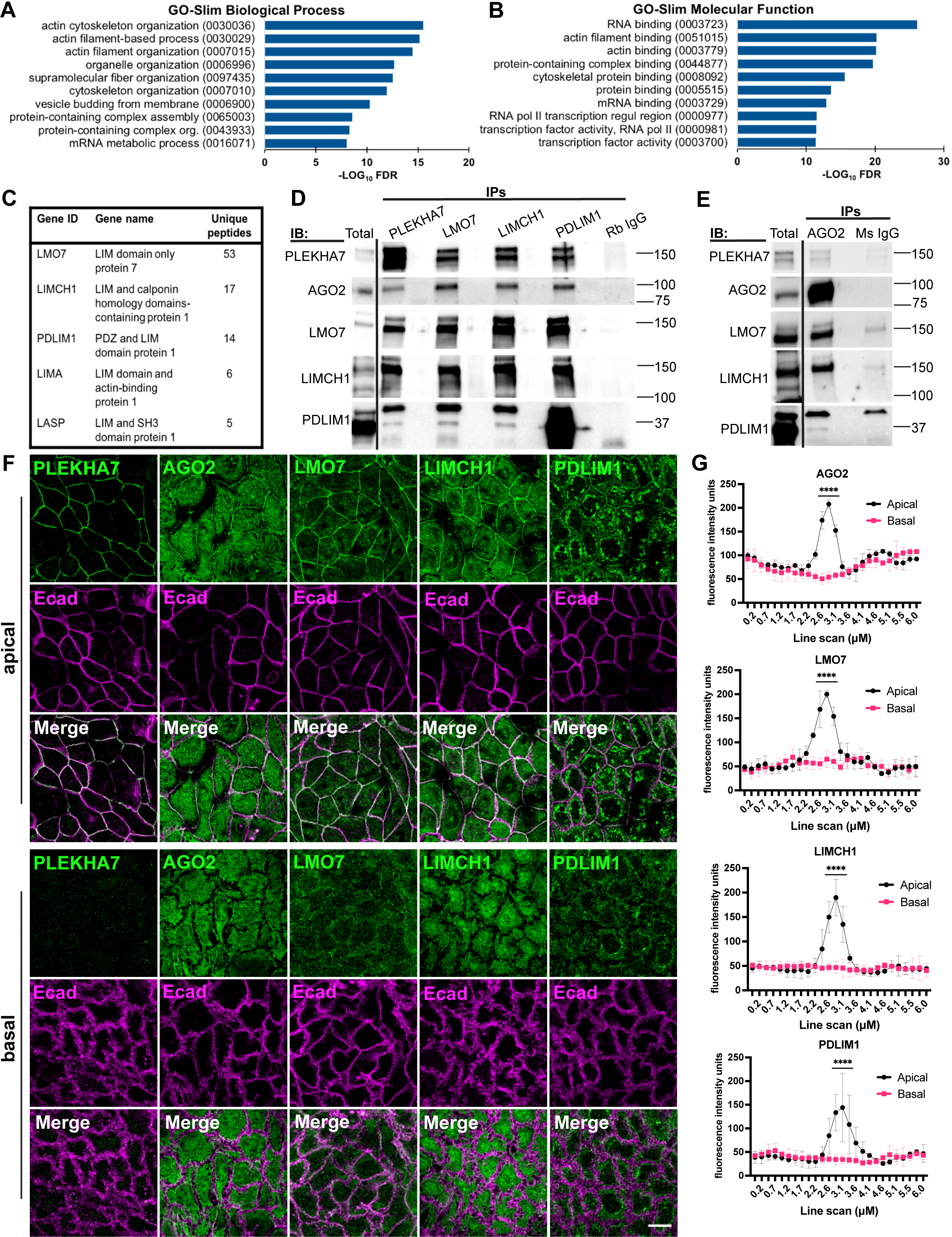
PLEKHA7 interacts with AGO2 and actin-related, LIM domain-containing proteins at apical adherens junctions. (A, B) Gene ontology (GO) enrichment analyses of the PLEKHA7-interacting protein dataset (http://www.imexconsortium.org/; identifier: IM-25739). The GO-slim Molecular Function and GO-slim Biological Process datasets were interrogated using Fisher’s exact test with false discovery rate (FDR) correction to assess enrichment significance. The 10 most significantly enriched GO terms are shown; GO term accession numbers are displayed in parentheses. (C) LIM domain-containing proteins identified in the same dataset of PLEKHA7 interactors; the unique peptide counts recorded from mass spectrometry analysis following PLEKHA7 immunoprecipitation are shown. (D, E) Immunoprecipitation (IP) of PLEKHA7, LMO7, LIMCH1, PDLIM1 and AGO2 from Caco2 cells, immunoblotted (IB) for the same markers; IgG is the negative control. Molecular masses (kD) are indicated on the right. (F-G) Immunofluorescence of E-cadherin (Ecad), PLEKHA7, LMO7, LIMCH1, PDLIM1 and AGO2 in confluent Caco2 cell monolayers. Images were obtained by confocal microscopy, and Z-series stacks were acquired through the entire plane; representative apical and basal Z-slices are shown. Fluorescence intensity of 6 µm line scans drawn perpendicular to cell-cell junctions was measured from n=30 cell-cell junctions (10 junctions/field) representative of three independent experiments; statistical analyses were performed using two-way ANOVA tests; error bars represent mean ± SD; ****, P < 0.0001. Scale bar = 20 µm.

Through examining the group of actin-related PLEKHA7 interactors, we noted several proteins belonging to the LIM domain-containing protein family (Figure 1C). These proteins attracted our attention, since LIM protein family members, such as Ajuba, LIMD1, and WTIP, have been previously implicated in AGO2 recruitment and RISC assembly in cytoplasmic p-bodies (Bridge et al., 2017; James et al., 2010; Tilley et al., 2020). Our PLEKHA7 proteomics analysis identified a different set of proteins belonging to this same family (Figure 1C). More specifically, the LIM domain protein with the highest count of PLEKHA7-enriched unique peptides was LIM domain only protein 7 (LMO7) (Figure 1C). LMO7 was previously identified to localize to apical adherens junctions, where it plays an actin-mediated role in linking and stabilizing nectin and cadherin junctional complexes (Ooshio et al., 2004). The other top enriched LIM domain-containing proteins were LIM and calponin homology domains-containing protein 1 (LIMCH1) and PDZ and LIM domain protein 1 (PDLIM1) (Figure 1C). Considering the reported functions of LIM domain-containing proteins in actin (Anderson et al., 2021; Kadrmas and Beckerle, 2004) and AGO2 regulation (Bridge et al., 2017; James et al., 2010; Tilley et al., 2020), we sought to interrogate these three LIM proteins as intermediates of AGO2 recruitment to the ZA. Protein co-immunoprecipitation, followed by Western blot analysis, confirmed PLEKHA7-LMO7-LIMCH1-PDLIM1 protein interactions with each other and with AGO2 (Figure 1, D and E). Furthermore, immunofluorescence and confocal microscopy analysis of Caco2 monolayers revealed localization of all three LMO7, LIMCH1, and PDLIM1 distinctly to the apical ZA, similar to AGO2 localization, as determined by co-localization with mature E-cadherin adhesions and the ZA-specific marker PLEKHA7 (Kourtidis et al., 2017b; Kourtidis et al., 2015) (Figure 1, F and G). Conversely, neither of these LIM proteins, AGO2, nor PLEKHA7, localize at basolateral areas of cell-cell contact (Figure 1, F and G). Together, these data point to the ZA as the subcellular compartment where PLEKHA7, AGO2, LMO7, LIMCH1, and PDLIM1 co-localize and interact; we also identify LIMCH1 and PDLIM1 as new ZA-associated LIM protein family members.

### PLEKHA7, LMO7, LIMCH1, and PDLIM1 are required for recruitment of AGO2 to the ZA

We have shown that AGO2 localization at the ZA depends on PLEKHA7 (Kourtidis et al., 2017b). Since LMO7, LIMCH1, and PDLIM1 interact with PLEKHA7 and AGO2 and all these proteins co-localize at the ZA, we asked whether any of these LIM proteins mediate recruitment of AGO2 to the ZA. Strikingly, shRNA-mediated knockdown of each of these LIM proteins resulted in loss of junctional AGO2 localization, without affecting AGO2 overall levels or cytoplasmic distribution, similarly to the effects of CRISPR/Cas9-generated PLEKHA7 knockout (Figure 2, A-D) and of our published PLEKHA7 shRNA-mediated knockdown (Kourtidis et al., 2017b). We also obtained identical results on AGO2’s junctional localization in PLEKHA7, LMO7, LIMCH1, or PDLIM1 - depleted cells, by using an exogenously introduced Flag-tagged AGO2 construct (AGO2-Flag; Figure S1, A-D), as well as by using a second, independent, set of LMO7, LIMCH1, PDLIM1 shRNAs (Figure S1, E-G). Furthermore, RNA immunoprecipitation and subsequent qRT-PCR analysis showed that LMO7, LIMCH1, or PDLIM1 knockdown resulted in decreased AGO2 loading of a set of miRNAs, namely miR-24, miR-200c, miR-203a (Figure 2E). We have previously shown that PLEKHA7 depletion results in decreased AGO2 loading of the same set of miRNAs, which is an indicator of decreased AGO2 activity (Kourtidis et al., 2017b). Interestingly, knockdown of neither LMO7, LIMCH1, nor PDLIM1 had any effect on the localization or overall cellular levels of PLEKHA7 (Figure 2, C and D), suggesting that LIM protein-mediated effects on AGO2 are downstream of PLEKHA7. Based on these findings, we hypothesized that PLEKHA7 knockout would result in loss of junctional localization of these proteins, leading to loss of junctional AGO2. Intriguingly, junctional localization of neither LMO7, LIMCH1, nor of PDLIM1 are affected by PLEKHA7 loss (Figure 3A). Similarly, in each of the three LIM protein knockdowns, junctional localization of the other two LIM proteins was also not affected (Figure 3, B-D), although knockdowns of all three LMO7, LIMCH1, and PDLIM1 result in loss of junctional AGO2. Taken together, these results indicate that although PLEKHA7, LMO7, LIMCH1, and PDLIM1 form a complex at the ZA (Figure 1, D-G) and are each individually required for recruitment of AGO2 to the ZA, their impact on AGO2 recruitment is independent of their complex formation and junctional recruitment. This suggests the existence of a downstream effector linked to all these proteins that regulates AGO2 recruitment to the junctions, which we sought to identify next.

**FIGURE 2.**
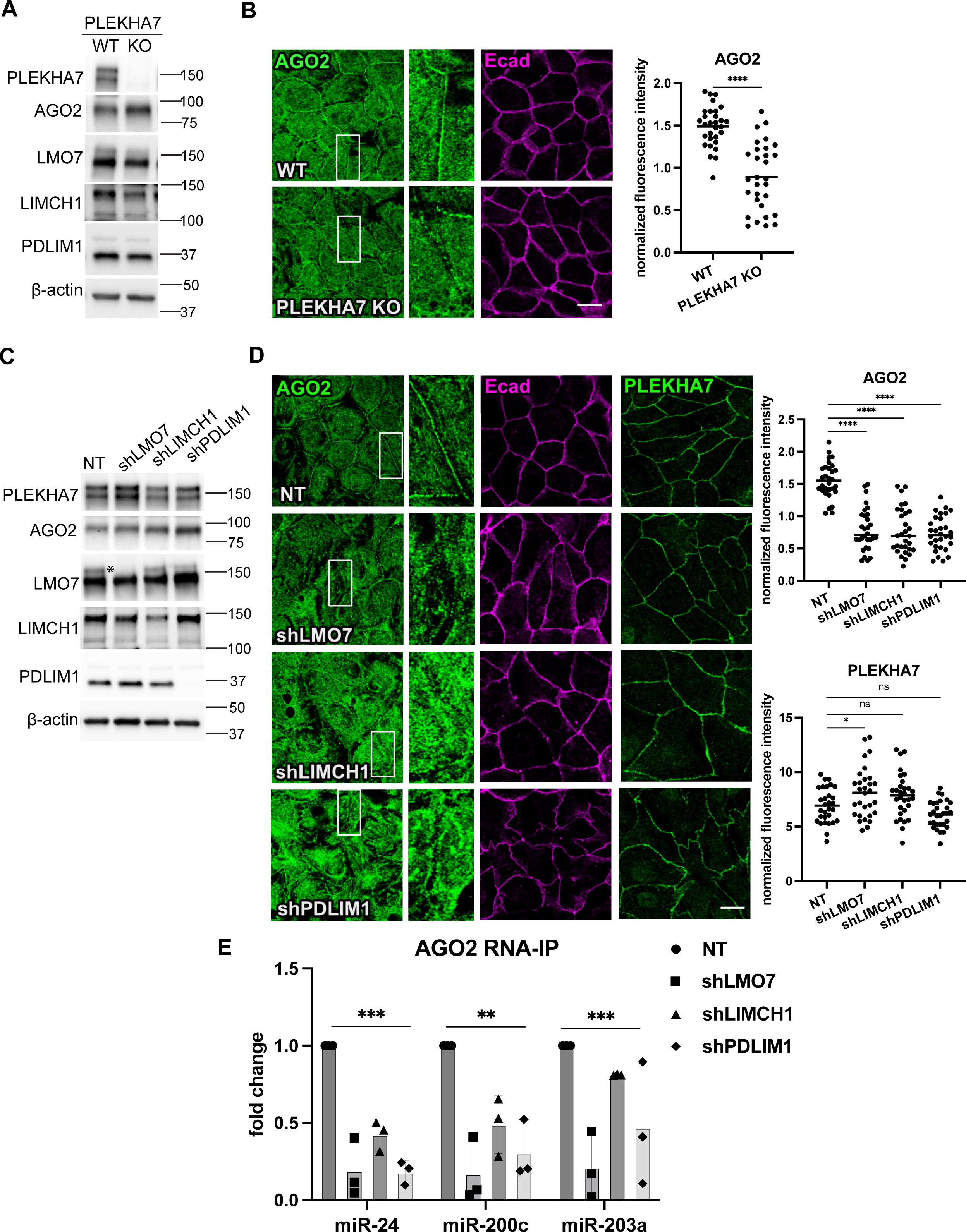
LMO7, LIMCH1, PDLIM1, and PLEKHA7 loss each disrupt junctional localization of AGO2. (A) Immunoblotting of PLEKHA7 knockout (KO) Caco2 cells, compared to control wild type cells (WT). (B) Immunofluorescence of control (WT) or PLEKHA7 knockout (KO) cells for AGO2 and E-cadherin (Ecad). AGO2 junctional fluorescence intensity normalized to cytoplasmic was quantified from n=30 cell-cell junctions (10 junctions/field) representative of three independent experiments; statistical analyses were performed using unpaired two-way *t* test; ****, P < 0.0001. (C) Immunoblotting of LMO7, LIMCH1, and PDLIM1 shRNA-mediated knockdown (shLMO7, shLIMCH1, shPDLIM1, respectively) Caco2 cells, compared to non-target (NT) shRNA control cells; asterisk indicates the specific LMO7 band lost by shRNA targeting. (D) Immunofluorescence of NT, shLMO7, shLIMCH1, shPDLIM1 Caco2 cells for AGO2, PLEKHA7, and E-cadherin (Ecad). AGO2 and PLEKHA7 junctional fluorescence intensity normalized to cytoplasmic was quantified from n=30 cell-cell junctions (10 junctions/field) representative of three independent experiments; statistical analyses were performed using one-way ANOVA test; ****, P < 0.0001; *, P < 0.05; ns, non-significant. Insets in all cases are marked by white rectangles and are 3X magnification of the original image. In all immunoblots, β-actin is the loading control; molecular masses (kD) are indicated on the right. Top-view immunofluorescence images were obtained by confocal microscopy and are single apical Z-slices. Scale bars = 20 µm. (E) AGO2 RNA immunoprecipitation (RNA IP) followed by qRT-PCR analysis for miR-24, miR-200c, and miR-203a miRNAs in NT, shLMO7, shLIMCH1, shPDLIM1 Caco2 cells. Error bars represent mean ± SD from n=3 independent experiments; statistical analysis was performed using one-way ANOVA ***, P < 0.005; **, P < 0.01.

**FIGURE 3.**
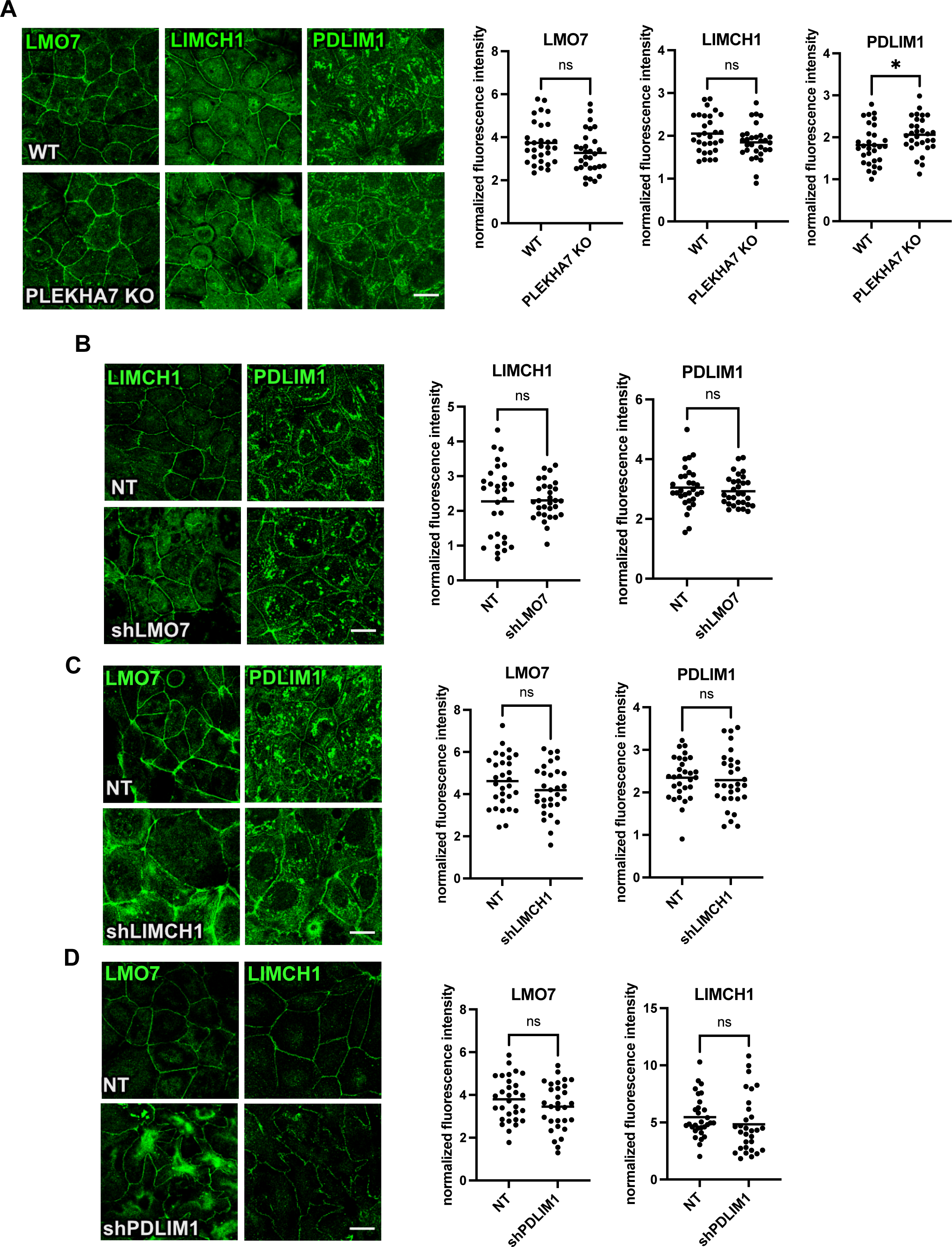
PLEKHA7, LMO7, LIMCH1, or PDLIM1 loss does not affect each other’s localization to the junctions. (A) Immunofluorescence of control (WT) or PLEKHA7 knockout (KO) cells for LMO7, LIMCH1, and PDLIM1. (B-D) Immunofluorescence of each LMO7, LIMCH1, and PDLIM1 in control (NT) and shLMO7, shLIMCH1,and shPDLIM1 Caco2 cells. For all figures, top-view immunofluorescence images were obtained by confocal microscopy and are single apical Z-slices. Junctional fluorescence intensity normalized to cytoplasmic was quantified from n=30 cell-cell junctions (10 junctions/field) representative of three independent experiments; statistical analyses were performed using unpaired two-way *t* test; *, P < 0.05; ns, non-significant. Scale bars = 20 µm.

### PLEKHA7, LMO7, LIMCH1, or PDLIM1 depletion disrupts actomyosin organization at the ZA

LMO7, LIMCH1, and PDLIM1 have been previously reported to be involved in actomyosin organization (Bauer et al., 2000; Lin et al., 2017; Maeda et al., 2009; Ooshio et al., 2004; Tamura et al., 2007; Vallenius et al., 2000). We have also shown that PLEKHA7 knockdown disrupts the integrity of the apical circumferential actin ring (Kourtidis et al., 2015), and our GO analysis reveals actin-related functions are significantly enriched in the PLEKHA7-interacting proteome (Figure 1, A and B). Based on these observations, we asked whether the relationship of these components with the actin cytoskeleton could be the common denominator regulating AGO2 recruitment to the ZA. First, we examined potential effects of these proteins on actomyosin organization in Caco2 cells specifically at the ZA, where AGO2 is recruited. Using confocal and super resolution microscopy we found that PLEKHA7 knockout cells exhibit disrupted actomyosin organization at the ZA, manifested by a disorganized circumferential actin ring and diffused junctional myosin IIB (Figures 4A and S2A). Similarly, knockdown of each of LMO7, LIMCH1, and PDLIM1 had distinct, but in all cases significant effects on actomyosin organization throughout the cell and particularly at the ZA (Figures 4B and S2, B and C). More specifically, in LMO7 knockdown cells, apical actin filaments failed to tightly bundle into a dense cable as in control cells but remained loosely organized in dispersed, multifurcated, actin cables (Figure 4B). LIMCH1 knockdown cells showed a different pattern, where the apical filamentous actin ring is severely fragmented or completely absent (Figure 4B). Finally, PDLIM1 knockdown cells exhibited jagged actomyosin cables alongside bicellular junctions that appeared to be attached to intracellular actomyosin cables linked to dense cytoplasmic actomyosin aggregates (Figure 4B - bottom panel). This further manifested in wavy junctional cell borders and with the appearance of a distorted, collapsed apical ring structure (Figure 4B). Intense stress fibers were also observed in LMO7, LIMCH1 and PDLIM1 knockdown cells (Figure S2B - basal panel). We particularly examined myosin IIB to assess actomyosin organization, since this is the non-muscle myosin II isoform specifically responsible for contractile tension and actin filament stabilization at the apical actin ring at the ZA (Gomez et al., 2015; Heuze et al., 2019; Smutny et al., 2010; Wayt et al., 2021). Indeed, effects on myosin IIA were not as consistent: although depletion of LMO7 and LIMCH1 resulted in disorganized and more diffused myosin IIA, they also resulted in stronger myosin IIA recruitment around areas of cell-cell contact, whereas depletion of PLEKHA7 or of PDLIM1 only minimally affected myosin IIA recruitment at the ZA (Figure S3, A and B). These results demonstrate that the negative effects of PLEKHA7, LMO7, LIMCH1, and PDLIM1 depletion on actomyosin organization at the ZA primarily involve myosin IIB. Altogether, these results show that PLEKHA7, as well as LMO7, LIMCH1, and PDLIM1, are each distinctly responsible for the stability and homeostasis of the actomyosin cytoskeleton and particularly of the circumferential actin ring at the ZA.

**FIGURE 4.**
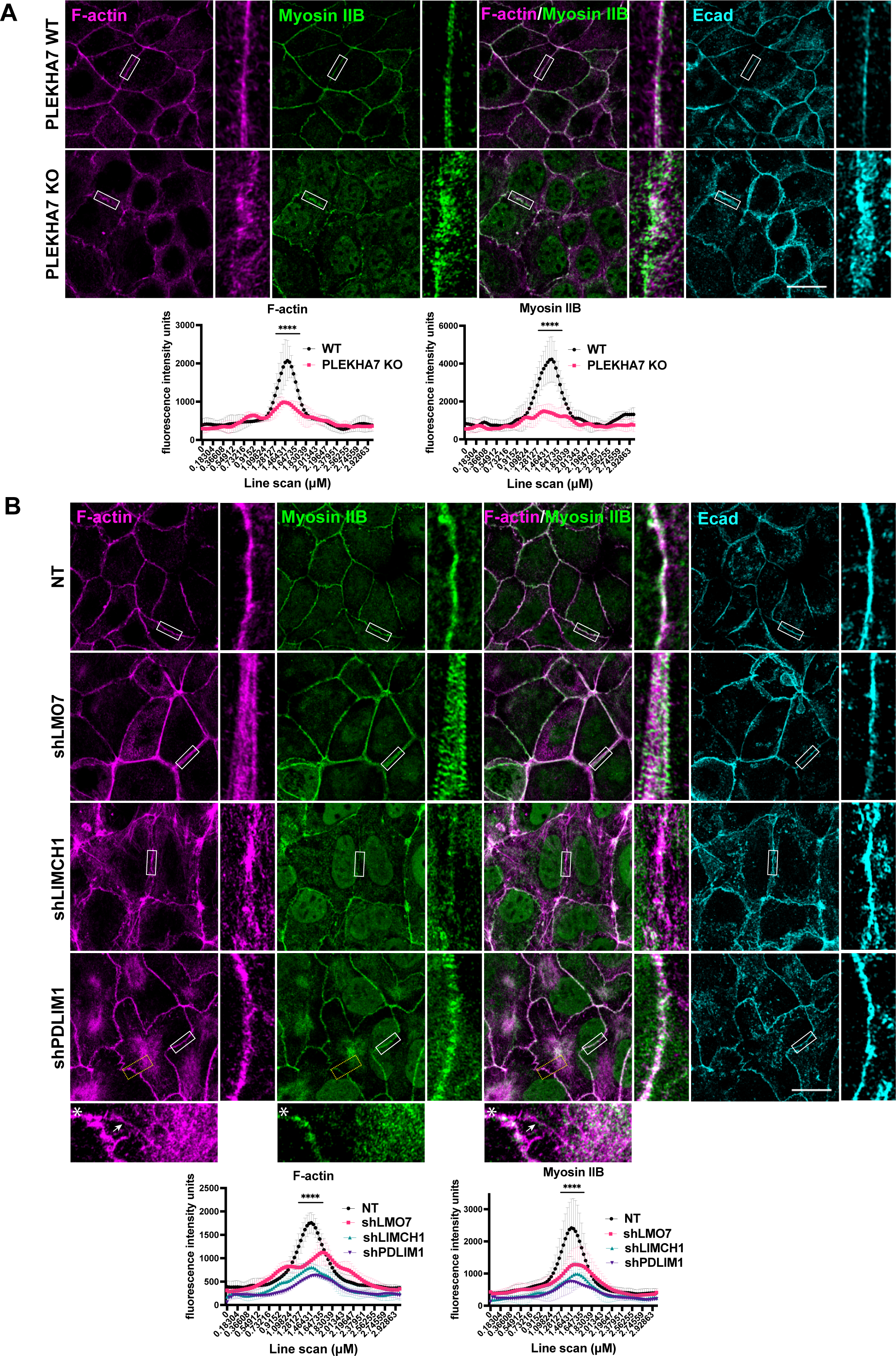
PLEKHA7, LMO7, LIMCH1, and PDLIM1 each distinctly influence apical actomyosin organization. (A) Immunofluorescence and super resolution microscopy of PLEKHA7 knockout (KO) Caco2 cells compared to control wild type (WT) Caco2 cells. (B) Immunofluorescence and super resolution microscopy of LMO7, LIMCH1, and PDLIM1 knockdown (shLMO7, shLIMCH1, shPDLIM1) Caco2 cells, compared to (NT) shRNA control cells. Top-view immunofluorescence images are maximum intensity projections of the two most apical Z-slices. Insets marked by white rectangles are 6X magnification of the original image. Insets for shPDLIM1 cells noted with asterisks are magnified 5X from areas marked with yellow dashed boxes. Asterisk marked insets are from full maximum intensity projection images to indicate basolateral actin filaments linked to the apical actin ring (marked by white arrows). In all cases, E-cadherin (Ecad) is used as a co-stain denoting adherens junctions. Fluorescence intensity of 3 µm line scans drawn perpendicular to cell-cell junctions was measured from n=30 cell-cell junctions (10 junctions/field) representative of three independent experiments; statistical analyses were performed using two-way ANOVA tests; error bars represent mean ± SD; ****, P < 0.0001. Scale bars = 20 µm.

### AGO2 recruitment to the ZA depends on the presence of an organized actin cytoskeleton

Since PLEKHA7, LMO7, LIMCH1, and PDLIM1 are all required for stability of the apical actomyosin cytoskeleton and for recruitment of AGO2 to the ZA, independently of each other, we asked whether proper actin organization is the underlying mechanism mediating recruitment of AGO2 to the ZA. To examine this, we utilized a classical calcium-switch assay (Gumbiner and Simons, 1986) to dismantle the calcium-dependent E-cadherin-based junctions and allow them to recover with or without the presence of Latrunculin A (LatA), an actin polymerization inhibitor (Coue et al., 1987) (Figure 5A). E-cadherin-based junctions and the junctional localization of PLEKHA7 and AGO2, although entirely lost upon calcium depletion, fully recover within 1 hour of calcium re-introduction in control cells (Figure 5, B-D). We also observe the same recovery time for the formation of a mature circumferential actin ring in these control cells (Figure 5A). These cells are entirely rounded before recovery begins, due to loss of cell-cell adhesion and actin contraction, which is marked by cytoplasmic, perinuclear actin rings (Figure 5A). However, as previously reported (Ivanov et al., 2004; Yu-Kemp et al., 2022), LatA-treated cells fail to round following calcium depletion, due to the lack of contraction, and pools of E-cadherin, as well as punctate actin pools, are retained at areas of cell-cell contact (Abe and Takeichi, 2008; Cavey et al., 2008; Tang and Brieher, 2012; Yamada et al., 2004) (Figure 5, A and D). Furthermore, E-cadherin re-populates areas of cell-cell contact to re-form adherens junctions during recovery in the LatA cells, although these cells do not contain filamentous actin, again in agreement with previous observations (Ivanov et al., 2004; Tang and Brieher, 2012; Yu-Kemp et al., 2022) (Figure 5, A and D). Interestingly, we also found that PLEKHA7 follows a similar pattern as E-cadherin in the LatA cells, mostly recovering at the junctions within 1 hour of calcium re-introduction (Figure 5C). Therefore, although PLEKHA7 is critical for the stability of the actin cytoskeleton, its localization to the junctions does not depend on the presence of organized, filamentous actin. Notably, pools of LMO7, LIMCH1, and of PDLIM1 are also retained at cell-cell junctions upon recovery in the LatA-treated cells (Figure 5, E-H). However, localization of AGO2 to the junctions is completely abolished in LatA-treated cells and never recovers after calcium re-addition (Figure 5, B and H). We obtained identical results using the AGO2-Flag construct (Figure S4A). These data, together with our observed effects on actin organization and AGO2 ZA recruitment by each one of PLEKHA7, LMO7, LIMCH1, and PDLIM1, demonstrate that AGO2 recruitment to the ZA depends on the presence of an organized, mature apical actin ring, downstream of all four PLEKHA7, LMO7, LIMCH1, and PDLIM1.

**FIGURE 5.**
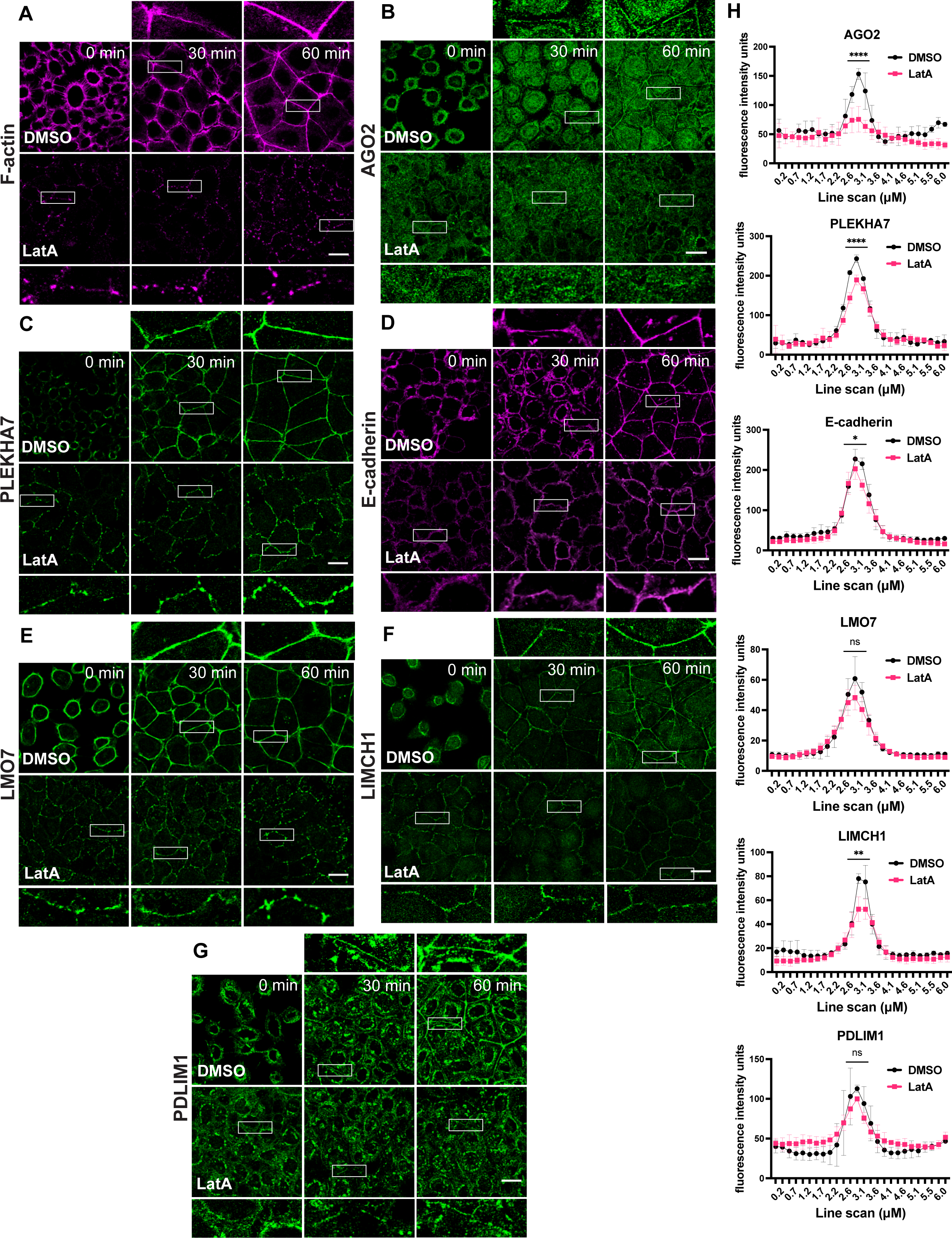
AGO2 recruitment to the ZA depends on filamentous actin integrity. (A-G) Immunofluorescence of Caco2 cells for F-actin, PLEKHA7, AGO2, E-cadherin, LMO7, LIMCH1, PDLIM1, fixed at indicated time points throughout a calcium switch assay. Images at 0 min indicate Ca^2+^ depleted cells immediately before Ca^2+^ reintroduction and at 30, 60 min time points are following Ca^2+^ re-addition. Cells were pre-treated and maintained in either DMSO vehicle (control) or 10 µM Latrunculin A (LatA) throughout the calcium switch assay. Insets are marked by white rectangles and are 3X magnification of the original image. Scale bars = 20 µm. (H) Fluorescence intensity of 6 µm line scans drawn perpendicular to cell-cell junctions was measured for each marker at the 60 min time point upon recovery from n=30 cell-cell junctions (10 junctions/field) representative of three independent experiments; statistical analyses were performed using two-way ANOVA tests; error bars represent mean ± SD; ****, P < 0.0001; **, P < 0.01; *, P < 0.05; ns, non-significant.

### PLEKHA7, LMO7, LIMCH1, or PDLIM1 depletion disrupts mechanical tension at the ZA

Our findings demonstrate that AGO2 recruitment to the ZA depends on the presence of a mature circumferential actin ring and on PLEKHA7, LMO7, LIMCH1, PDLIM1, which are all required for the stability of the actin cytoskeleton. However, we also noted that depleting each of these components had varied effects on apical actomyosin; apical actin filaments were present but dispersed in LMO7 and PDLIM1 knockdown cells (Figure 4B), whereas apical actin was fragmented and disorganized in LIMCH1 knockdown and PLEKHA7 knockout cells (Figure 4, A and B). In most cases, apical actin was also notably dissociated from myosin. This is informative, since myosin monomers assemble to form an ATPase-powered motor that imposes contractile forces on the AJ-associated actin cytoskeleton (Vicente-Manzanares et al., 2009). The dissociation of apical actin and myosin in the LIM knockdown and in PLEKHA7 knockout cells, points to decreased F-actin mechanical tension (Uyeda et al., 2011). To evaluate the effects of PLEKHA7, LMO7, LIMCH1, and PDLIM1 on contractile tension of the apical actin ring, we used the tension-sensitive α18 antibody, which recognizes an α-catenin epitope that becomes exposed when structural changes are induced by actomyosin tension (Yonemura et al., 2010). Indeed, depletion of each of PLEKHA7, LMO7, LIMCH1, and PDLIM1 disrupted actomyosin tension at the ZA, as indicated by decreased α18 intensity at bicellular junctions, although total junctional α-catenin remained unaffected (Figure 6, A - D). Notably, each of these cell populations display distinct cell shape changes, further reflecting disruption of tension distribution (Levayer and Lecuit, 2012). More specifically: a) LMO7 knockdown cells are uneven in size, with either very large or small cells observed; b) LIMCH1 knockdown cells are uniformly large and flattened; and c) PDLIM1 knockdown cells are smaller with distinct, wavy apical cell-cell borders (Figure 6C). Wavy apical cell-cell junctions, or otherwise loss of junction straightness, has also been previously associated with reduced contractile cortical tension on the junctions (Arnold et al., 2019; Sumi et al., 2018; Yamamoto et al., 2021). Indeed, measurements of bicellular junction lengths show loss of junctional straightness in PDLIM1 knockdown cells (Figure 6E). Along these lines, depletion of each of PLEKHA7, LMO7, LIMCH1, and PDLIM1 also resulted in decreased phosphorylation of myosin II regulatory light chain at Serine 19 (pS19-MRLC), as well as in its uneven distribution along cell-cell junctions, further indicating disruption of actomyosin tension (Figure 6, F-I). Altogether, these results demonstrate that PLEKHA7, LMO7, LIMCH1, and PDLIM1 are all responsible for the establishment and proper distribution of tension at the ZA.

**FIGURE 6.**
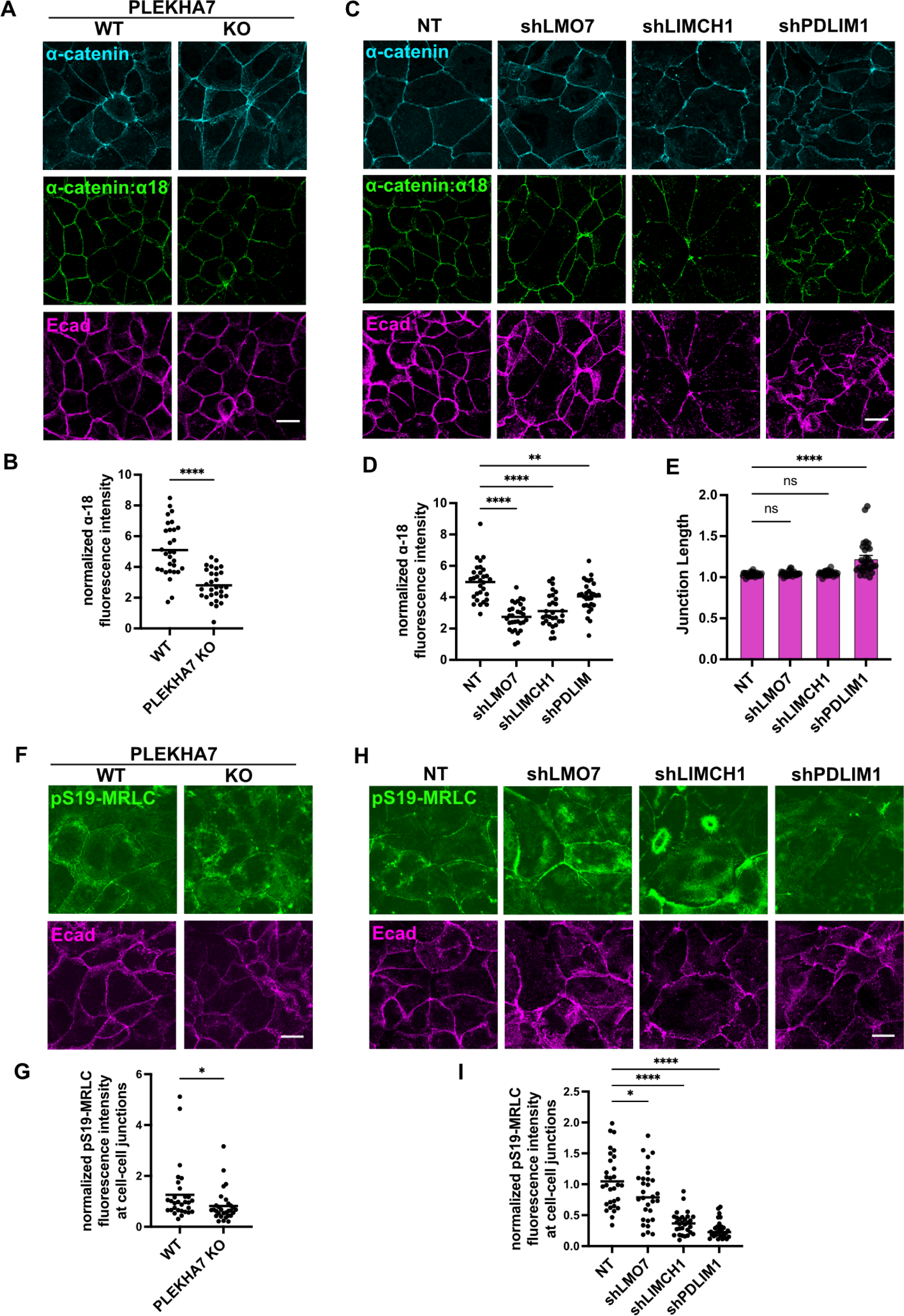
Contractile tension at the ZA is disrupted in LMO7, LIMCH1, PDLIM1 knockdown and PLEKHA7 knockout cells. Immunofluorescence of (A-E) total, as well as of tensile α-catenin using the α18 antibody that recognizes α-catenin upon tension, as well as of (F-I) phosphorylated myosin light chain at S19 (pS19-MRLC) that also indicates tension, in wild type (WT) or PLEKHA7 knockout (KO) Caco2 cells and in control (NT) or LMO7, LIMCH1, and PDLIM1 knockdown (shLMO7, shLIMCH1, shPDLIM1) cells. E-cadherin (Ecad) is used as a co-stain denoting adherens junctions. α18 fluorescence intensity measurements were normalized to total α-catenin (B, D) and pS19-MRLC were normalized to E-cadherin (G, I). All quantifications are from n=30 cell-cell junctions (10 junctions/field) representative of three independent experiments. Statistical analyses were performed using unpaired two-way *t* tests (B, G) or one-way ANOVA tests (D, E, I). ****, P < 0.0001; **, P < 0.01; *, P < 0.05; ns, non-significant. Scale bars= 20 µm.

### AGO2 recruitment to the ZA depends on actin tension and interaction with Myosin IIB at the ZA

We next asked whether contractile actomyosin tension at the ZA is also involved in the recruitment of AGO2 to the ZA. To examine this, we treated cells with: a) Blebbistatin, which inhibits the ATPase-dependent motor activity of myosin, but does not prevent its binding to actin (Kovacs et al., 2004); or b) the ROCK inhibitor, Y-27632, which indirectly inhibits myosin II motor activity by preventing MRLC phosphorylation (Amano et al., 2010). We performed these treatments during recovery, after calcium switch. Both Blebbistatin and Y-27632 resulted in decreased cortical actin ring tension, as indicated by disrupted actomyosin filaments and decreased α18 signals at the ZA (Figure S4, B-D). Concomitantly, AGO2 junctional localization was also disrupted in cells with reduced contractile tension, when using both Blebbistatin and Y-27632, and as demonstrated by: a) loss of endogenous AGO2 localization to the ZA; b) loss of endogenous AGO2 co-localization to α18; c) loss of ZA localization of the ectopically expressed AGO2-Flag construct (Figures 7A; S4, E and F; and S5). Altogether, these results demonstrate that AGO2 localization at the ZA is sensitive to disruption of actin tension.

**FIGURE 7.**
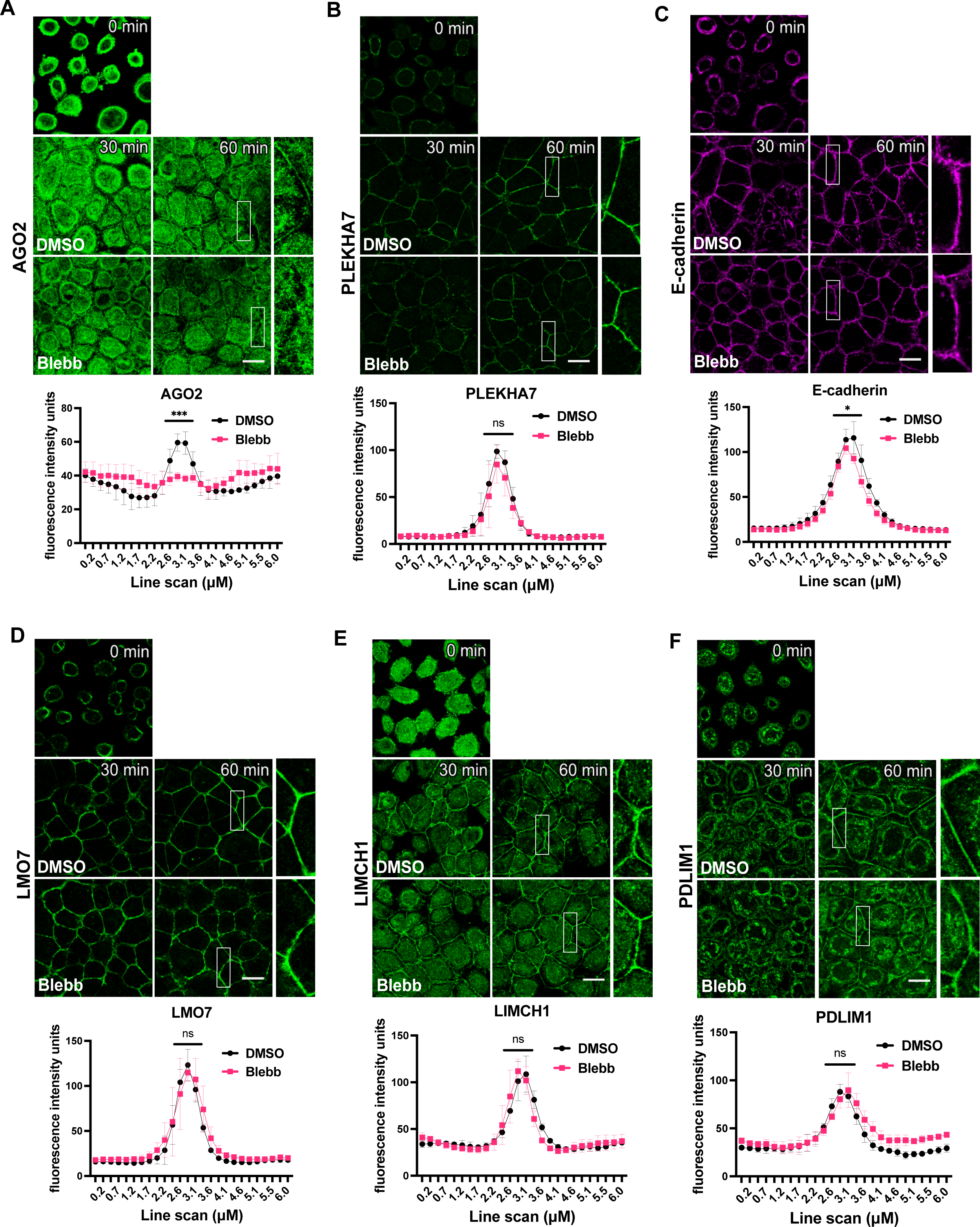
Contractile actin tension at the ZA is required for AGO2 junctional recruitment. (A-F) Immunofluorescence of Caco2 cells during a calcium switch assay in which DMSO or Blebbistatin were included in the calcium-containing recovery medium. Cells fixed at 30, 60 min post Ca^2+^ reintroduction were stained for AGO2 (A), PLEKHA7 (B), E-cadherin (C), LMO7 (D), LIMCH1 (E), and PDLIM1 (F). Insets are marked by white rectangles and are 3X magnification of the original image. Fluorescence intensity of 6 µm line scans drawn perpendicular to cell-cell junctions was measured for each marker at the 60 min time point upon recovery from n=30 cell-cell junctions (10 junctions/field) representative of three independent experiments; statistical analyses were performed using two-way ANOVA tests. Error bars represent mean ± SD. ****, P < 0.0001; *, P < 0.05; ns, non-significant. Scale bars= 20 µm.

In contrast, PLEKHA7, LMO7, LIMCH1 and PDLIM1 localization to the ZA were unaffected by either Blebbistatin, or ROCK inhibition (Figures 7, B-F; and S5, B-F). Taken together with the LatA findings (Figure 5), and the PLEKHA7, LMO7, LIMCH1, PDLIM1 depletion experiments (Figures 2 and 3), these results further demonstrate that the effects of PLEKHA7, LMO7, LIMCH1 or PDLIM1 depletion on AGO2 recruitment at the ZA are independent of either their junctional recruitment, or their protein-protein interactions, but rather associated with their downstream effects on actomyosin organization and tension. These results also suggest direct interaction of AGO2 with actomyosin. Since Myosin IIB was the actomyosin motor uniformly disrupted in all cases of PLEKHA7, LMO7, LIMCH1 and PDLIM1 depletion, we examined whether interaction with Myosin IIB mediates AGO2 localization at the ZA. Indeed, co-immunoprecipitation analysis revealed that AGO2 interacts with Myosin IIB (Figure 8A). We then implemented a proximity ligation assay (PLA) to assess whether this AGO2-Myosin IIB interaction particularly occurs at the ZA. We used the ectopically expressed AGO2-Flag construct for this assay to avoid any non-specific interactions with endogenous proteins and since we showed that this construct mimics the endogenous AGO2 by: a) identically localizing at the ZA; and b) this localization being similarly lost by LatA and Blebbistatin treatments in calcium switch assays (Figures S1A; S4, A and F). Indeed, the AGO2-Flag:Myosin IIB PLA assay revealed interactions of AGO2 with Myosin IIB both at the ZA and the cytoplasm. However, the ZA-localized interactions were the ones that were specifically disrupted in all cases of PLEKHA7, LMO7, LIMCH1 and PDLIM1 depleted cells (Figure 8, B-E). In summary, our data demonstrate that actomyosin organization and tension at the ZA is the common downstream effector of all four PLEKHA7, LMO7, LIMCH1 and PDLIM1 in mediating AGO2 recruitment to the ZA, through regulating its interaction with Myosin IIB.

**FIGURE 8.**
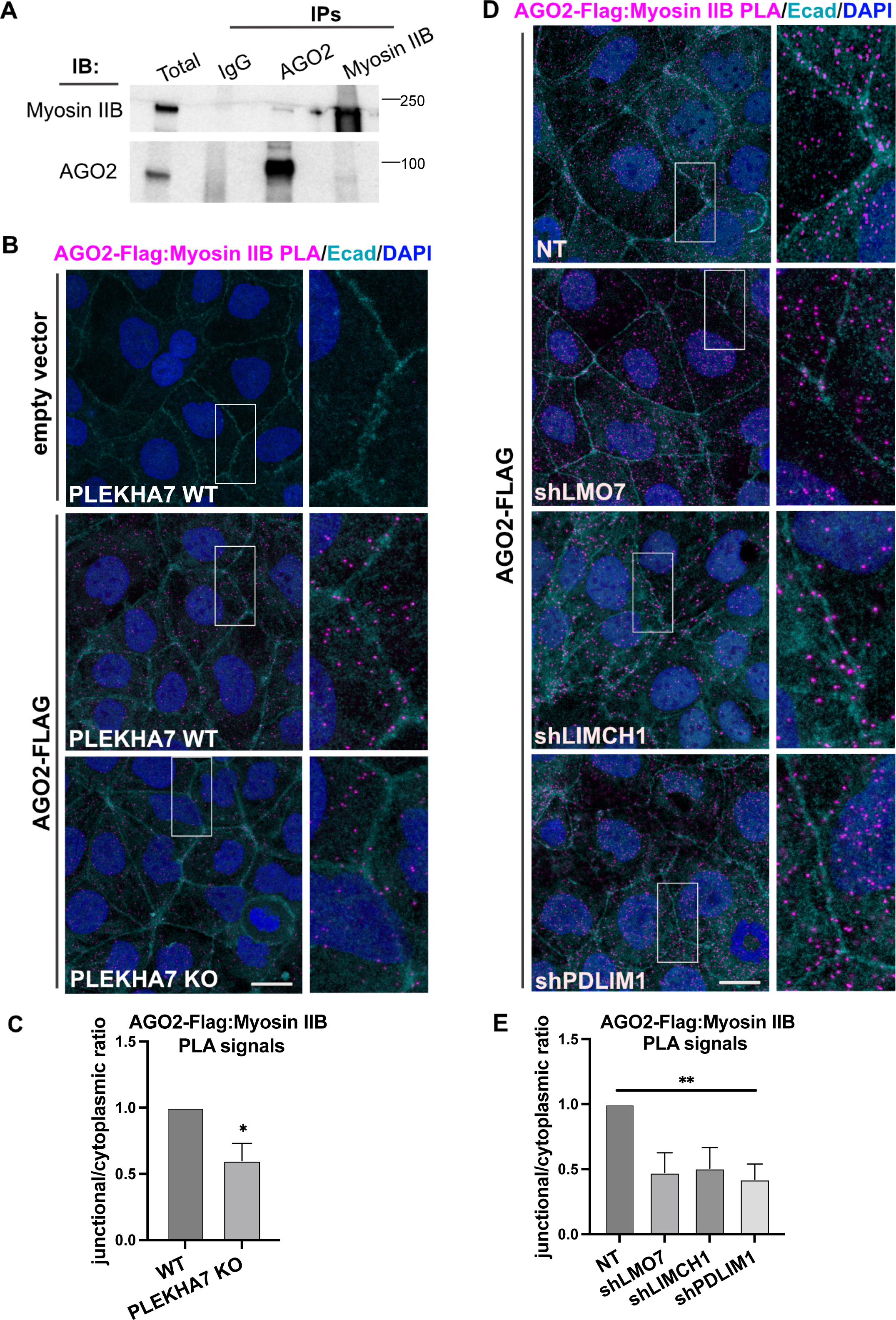
PLEKHA7, LMO7, LIMCH1, or PDLIM1 depletion each disrupt AGO2-Myosin IIB interaction at the ZA. (A) Immunoprecipitation (IP) of AGO2 and Myosin IIB from Caco2 cells, immunoblotted (IB) for the same markers; IgG is the negative control. Molecular masses (kD) are indicated on the right. (B-E) Wild type (WT) or PLEKHA7 knockout (KO) Caco2 cells and control (NT) or LMO7, LIMCH1, and PDLIM1 knockdown (shLMO7, shLIMCH1, shPDLIM1) cells stably expressing an AGO2-Flag construct (see Figure S1) were subjected to proximity ligation assay (PLA) for AGO2-Flag, using an anti-Flag antibody, and Myosin IIB, followed by immunofluorescence staining for E-cadherin (Ecad) and confocal microscopy; DAPI was used to stain nuclei. Caco2 cells stably transduced with an empty vector were used as negative PLA control. Insets are marked by white rectangles and are 3X magnification of the original image. The ratio of junctional vs cytoplasmic PLA signals was quantified for each condition and from n=30 cells (10 cells/field) representative of three independent experiments; statistical analyses were performed using either unpaired two-way *t* tests (B-C) or one-way ANOVA tests (D-E). Error bars represent mean ± SD. **, P < 0.01; *, P < 0.05. Scale bars = 20 µm.

## DISCUSSION

We have previously demonstrated that all major complexes of the RNAi machinery, namely the microprocessor complex, the DICER complex, as well as the RISC, including its main enzymatic component AGO2, distinctly localize at apical areas of cell-cell contact in well-differentiated epithelial cells and tissues and more specifically at the mature apical adherens junctions that form the ZA (Kourtidis et al., 2017b; Kourtidis et al., 2015; Nair-Menon et al., 2020). We have also shown that this ZA-associated RNAi machinery recruits and regulates processing and silencing activity of a set of miRNAs that suppress pro-tumorigenic phenotypes (Kourtidis et al., 2017b; Kourtidis et al., 2015). Moreover, we found that this ZA-specific localization is disrupted in tumors from patients (Kourtidis et al., 2015; Nair-Menon et al., 2020), further supporting the idea of a tumor-suppressing role of the ZA-associated RNAi machinery in cancer. However, a key fundamental question emerged from these findings: why are these complexes specifically present at the ZA of well-differentiated epithelial cells and tissues, to begin with?

To address this question, we set off in this work to better understand the mechanistic underpinnings of this junctional RNAi recruitment. In our previous work, we identified PLEKHA7 as the ZA-specific component responsible for recruitment of RNAi complexes to the junctions (Kourtidis et al., 2017b; Kourtidis et al., 2015; Nair-Menon et al., 2020). However, it remained unclear how PLEKHA7 recruits these RNAi components to the ZA. Through re-examining the PLEKHA7 proteome, we identified a set of LIM domain – containing proteins, namely LMO7, LIMCH1, PDLIM1, as novel PLEKHA7 binding partners at the ZA. These proteins attracted our attention, since other members of the same family of proteins have been identified in the past as responsible for recruitment of AGO2 to cytoplasmic p-bodies (Bridge et al., 2017; James et al., 2010; Tilley et al., 2020). Therefore, we examined the possibility that one of these proteins acts as the mediator between PLEKHA7 and AGO2 in recruiting it to the ZA. Surprisingly, we found that not only all three LMO7, LIMCH1, PDLIM1 are equally and individually required for recruitment of AGO2 at the ZA and of its miRNA loading activity, as PLEKHA7 is, but also that this recruitment is independent of their interactions, either between them or with PLEKHA7, at the ZA. Instead, what our experimentation revealed is that all four PLEKHA7, LMO7, LIMCH1, PDLIM1 are responsible for the stability of the actomyosin cytoskeleton at the ZA. We also found that this actomyosin stability, and more specifically the formation and maintenance of a mature, organized, tensile circumferential actin ring at the ZA, is the common denominator downstream of all four components that is responsible of AGO2 recruitment to the ZA. These findings have three major implications.

Firstly, they explain why AGO2 is specifically recruited to apical cadherin junctions at the ZA but not at basolateral areas of cell-cell contact, as we previously demonstrated both in cells (Kourtidis et al., 2017b) and in tissues (Nair-Menon et al., 2020) and further confirm in this study (Figure 1, F and G). Although we have demonstrated this localization so far in colon and kidney epithelial cells (Caco2; MDCK) (Kourtidis et al., 2017b) and in normal colon tissues (Nair-Menon et al., 2020), our current findings allow us to make the prediction that junctional AGO2 and its ensuing miRNA regulation may also be found in other epithelial cells and tissues that are well-differentiated and exhibit a mature, tensile circumferential apical actin ring and mature ZA. A similar prediction can be made not only when referring to human epithelia, but also to epithelia from other species, especially when considering that we have already seen this localization in at least one more species (canine MDCK cells). Since we have shown that PLEKHA7 is vertebrate-specific and has no invertebrate ortholog (Kourtidis et al., 2022), it would be particularly interesting to examine whether junctional AGO2 exists in invertebrate species, for example by virtue of LMO7, which has invertebrate orthologs (Beati et al., 2018), or other similar actin-binding components. It remains to be seen whether these predictions will be confirmed in future studies.

Secondly, our findings suggest that any perturbations of the circumferential actin ring, which can occur during normal processes, such as tissue morphogenesis and junctional remodeling, or aberrantly, during disease, can disrupt AGO2 localization and function. For example, it is known that aberrant actin remodeling is common in cancer (Hayward et al., 2021), and we have shown that junctional localization of AGO2 is broadly lost in colon tumors and cancer cell lines (Nair-Menon et al., 2020). These findings suggest that the mechanism instructing association of AGO2 to the ZA is much broader than any single protein-directed interactions and reveal an underlying cellular mode of regulation that is more fundamental in nature. Corroborating this notion, we show here that depletion of any one of PLEKHA7, LMO7, LIMCH1, and PDLIM1 leads to disruption of both structure and tension of the apical actomyosin ring and in loss of AGO2 junctional localization and miRNA-loading ability. Therefore, mutations or downregulation of any of these components, or of many more actin-regulating proteins, may feed into the same mechanism as a common central node of regulation, promoting disease progression.

Thirdly, this work provides a conceptual framework, which can guide us to further study this localized RNAi function and to fully realize its physiological significance, since we have shown that the ZA-localized RNAi machinery is critical for processing and function of a set of miRNAs that regulate cell behavior (Kourtidis et al., 2017b; Kourtidis et al., 2015; Nair-Menon et al., 2020). In this framework, RNAi complexes at the ZA act as sensors of actomyosin contractility and organization, directly linking the mechanical state of the cell with post-transcriptional regulation of gene expression and consequently rapid modulation of cell behavior. Although more experimentation is needed to fully establish this idea, these findings introduce the concept of “mechanosensitive RNAi”. In this report, we focus specifically on AGO2, however, we can also apply this approach to study the other RNA-binding proteins and complexes that we have identified at the ZA (Kourtidis et al., 2017b; Kourtidis et al., 2015) and to fully reconstruct their regulation and roles in cell and tissue homeostasis and disease. For example, we have recently shown that different extracellular matrix (ECM) components distinctly impact localization and complex formation of RNAi components to the ZA, including not only RISC and AGO2, but also DROSHA and the microprocessor (Daulagala and Kourtidis, 2022). It is well-established that different ECM components exert different mechanical forces in cells, which are primarily transmitted through the actomyosin cytoskeleton (Janmey et al., 2020). In light of our current findings, we can also make the prediction that these ECM-mediated effects on junctional recruitment and complex formation of RNAi complexes are indeed mediated through fine-tuning of the actin cytoskeleton, which remains to be tested in future studies.

Beyond its potential significance for the physiological relevance of the junctional AGO2 and RNAi, our findings may have additional repercussions on our understanding of the physiological roles of RNAi and miRNAs. For example, the actomyosin cytoskeleton has been implicated in localization and assembly of RNA-binding complexes in other cellular compartments, such as p-bodies in the cytoplasm or Cajal bodies and the RNA-induced transcriptional gene silencing-like (RITS-like) complex in the nucleus (Ahlenstiel et al., 2012; Lindsay and McCaffrey, 2011; McCaffrey and Lindsay, 2012; Skare et al., 2003). Furthermore, a more recent study showed that AGO2 in endothelial cells and fibroblasts engages miRNAs that particularly regulate biomechanical proteins and pathways, in response to being plated on stiff substrates (Moro et al., 2019). The above, taken together with our current findings, suggest that RNAi and miRNAs may indeed be a means for the cell to rapidly respond to biomechanical stimuli, further supporting the concept of “mechanosensitive RNAi” as a broader mechanism regulating cellular plasticity.

The present work also provides new information for the potential cellular functions of PLEKHA7, LMO7, LIMCH1, and PDLIM1. More specifically, we show that LMO7, LIMCH1, and PDLIM1 all co-exist as a complex at the apical ZA, together with PLEKHA7 and AGO2. Localization of LMO7 at apical AJs has been previously reported (Matsuda et al., 2022; Ooshio et al., 2004), but not that of LIMCH1 and PDLIM1, since previous studies primarily focused on their roles in localizing with and modulating stress fibers (Bauer et al., 2000; Lin et al., 2017; Maeda et al., 2009; Tamura et al., 2007; Vallenius et al., 2000). Then, we reveal that LMO7, LIMCH1, and PDLIM1 all regulate AGO2 recruitment to the ZA and miRNA loading to AGO2, identically to PLEKHA7 (Figure 2)(Kourtidis et al., 2017b). We also show that all four PLEKHA7, LMO7, LIMCH1, and PDLIM1 regulate actomyosin stability, albeit in seemingly different ways and potentially through different mechanisms. Using super-resolution microscopy, we show that PLEKHA7 depletion results in disruption of the actomyosin structure at the ZA, adding to our previously published findings showing that PLEKHA7 depletion results in cofilin activation and actin ring disruption (Kourtidis and Anastasiadis, 2016; Kourtidis et al., 2015). Furthermore, our PLEKHA7 proteomics analysis identified high enrichment in actin-binding proteins (Figure 1, A and B), which builds on our previous work that revealed association of PLEKHA7 with certain components of the actin cytoskeleton, such as ACTN1 and MYL6 (Kourtidis and Anastasiadis, 2016). A recent study also demonstrated that PLEKHA7 is linked to Myosin IIB through CGLN1 (Rouaud et al., 2023), which could also explain the effects of PLEKHA7 depletion on Myosin IIB distribution and actomyosin tension. We also show that LMO7 depletion negatively impacts actomyosin organization by resulting in a multifurcated apical actomyosin ring and extended, widened apical myosin arrays, an appearance previously described as “sarcomeric” (Choi et al., 2016; Yu-Kemp et al., 2022). This actomyosin organization is indicative of incomplete maturation of apical actin bundles (Choi et al., 2016; Yu-Kemp et al., 2022). In addition, LMO7-depleted junctions seem to be uneven in size and are less tensile. It has been shown that *Drosophila* mutants lacking expression of Smallish (Smash), the fly ortholog of LMO7, also showed junctions of unequal shape in the larval epidermis, whereas *Drosophila* Smash or *Xenopus* LMO7 overexpression resulted in increased apical constriction, further supporting a conserved role for LMO7 in promoting actomyosin contractility (Beati et al., 2018; Matsuda et al., 2022). We also observed actomyosin disruption, increased cell spreading and decreased contractility in LIMCH1 knockdown cells, similarly to what was previously reported in stress fibers in HeLa cells (Lin et al., 2017). Strikingly, our PDLIM1-depleted cells exhibited a notably distinct phenotype, with distorted cell borders that resemble those observed when the cortical actomyosin-associated protein Merlin is depleted in Caco2 cells (Chiasson-MacKenzie et al., 2015). This phenotype was associated with increased contraction of the medioapical cortical cytoskeleton and reorientation of pulling forces inward, perpendicular to cell-cell junctions (Chiasson-MacKenzie et al., 2015), contrasting contraction of the circumferential actomyosin belt, which exerts pulling forces parallel to the bicellular junctions. This loss of parallel junctional tension, coupled with increased medioapical contraction, has been associated with the presence of F-actin bundles linked perpendicular to the junctions (Arnold et al., 2019), which we also noted in PDLIM1 knockdown cells (Figure 4B). Altogether, these observations strongly support critical and distinct roles of LMO7, LIMCH1, and PDLIM1 on actomyosin stability at epithelial ZA. However, the molecular details of these roles remain to be thoroughly investigated in subsequent studies.

In conclusion, our study demonstrates that AGO2 recruitment and function at epithelial cell-cell junctions responds to the presence of a structurally mature and mechanically tensile actomyosin cytoskeleton at the ZA, introducing the concept of “mechanosensitive RNAi”. Our findings also suggest that this mechanosensitive RNAi may be a broader, more fundamental property of the cell, adding another layer of regulation of cellular plasticity, which can have implications in our understanding of developmental or diseased processes.

## MATERIALS AND METHODS

### Cell culture, constructs, generation of stable cell lines

In all comparisons, cells were used at strictly the same confluencies. All cell lines were authenticated by the University of Arizona Genetics Core (via Science Exchange) and checked for misidentified, cross-contaminated, or genetically drifted cells. Cell lines tested negative for mycoplasma contamination (LookOut Mycoplasma PCR Detection Kit, Sigma Aldrich). Caco2 colon epithelial cells (ATCC, #HTB-37) were cultured in MEM (Corning) supplemented with 10% FBS (Gibco), 1 mM sodium pyruvate (Gibco), and 1X nonessential amino-acids (NEAA) solution (Gibco). HEK 293FT human embryonic kidney cells (Thermo Fisher Scientific, #R70007) were cultured in DMEM (Corning) supplemented with heat-inactivated 10% FBS, 2 mM additional L-glutamine, 1 mM sodium pyruvate, and 1X NEAA. Cells were maintained at 37°C with 5% CO_2_. Lentiviral shRNA constructs were derived from the pLKO.1-based TRC1 shRNA library (Sigma-Aldrich/RNAi Consortium); the following vectors were used: pLKO.1-puro Non-Target shRNA Control, SHC016; shLMO7, TRCN0000006491; shLIMCH1, TRCN0000153799; shPDLIM1, TRCN0000161271; 2^nd^ set: shLMO7, TRCN0000006492; shLIMCH1, TRCN0000154280; shPDLIM1 TRCN0000163210. The pLJM1-FLAG-HA-AGO2-WT (Addgene plasmid # 91978; http://n2t.net/addgene:91978; RRID:Addgene_91978) and pLJM1-Empty (Addgene plasmid # 91980; http://n2t.net/addgene:91980; RRID:Addgene_91980) lentiviral vectors were a gift from Joshua Mendell (Golden et al., 2017). Lentiviral particles were produced in HEK 293FT cells co-transfected via Lipofectamine 2000 (Invitrogen) with the shRNA vector and Virapower packaging plasmids (pMDLg/pRRE, Addgene #12251; pRSV-Rev, Addgene #12253; pMD2.G, Addgene #12259) according to manufacturer instructions. shRNA lentivirus was collected and used to infect Caco2 cells according to standard protocols. Stable shRNA-expressing cell lines were maintained in 2.5 μg/ml puromycin (Gibco) following initial selection at 5 μg/ml.

The PLEKHA7 knockout Caco2 cell line was generated using CRISPR/Cas9 technology, as previously described (Ran et al., 2013). The sgRNA sequence (GGATGTTTCACTGACCATGCTGG) targeting exon 5 of PLEKHA7 was designed using the online Synthego tool (www.synthego.com) and synthesized by Integrated DNA Technologies, IDT. This PLEKHA7-targeting sgRNA was cloned into the PX459 pSPCas9(BB)-2A-Puro vector (Ran et al., 2013). Caco2 cells were transfected with 30 µg the sgRNA-containing vector using Lipofectamine 3000 (Invitrogen) according to the manufacturer’s protocol. After 24 hours, cells were selected with 5 μg/ml puromycin for 48 hrs. Successfully transfected cells were seeded to obtain single colonies, which were picked and expanded for screening. PCR using primers designed to amplify the target region (Forward: CCTTTCTCAGGGCCTCATGTCA; Reverse: GCCTCCAAACAATCAGGGTTGG) was performed with DNA extracted from colonies using QuickExtract (Lucigen). PCR amplified products were screened by restriction digest analysis using the CviAII restriction enzyme (New England Biolabs). Positive candidates were sequenced to identify and confirm indels (Figure S6).

### Antibodies

The primary antibodies used in this study were PLEKHA7 (HPA038610; Sigma-Aldrich), E-cadherin (610182; BD Transduction Labs), AGO2 (ab57113; Abcam - optimal for immunoblotting), AGO2 (AP5281; ECM Biosciences - optimal for immunofluorescence), AGO2 (ab156870; Abcam - optimal for immunofluorescence and immunoprecipitation/RNA-immunoprecipitation); LMO7 (ab224113; Abcam), LIMCH1 (ab96178; Abcam), PDLIM1 (ab129015; Abcam), β-actin (4967L; Cell Signaling Technology), Myosin IIB (909901; Biolegend), Myosin IIA (909801; Biolegend), pS19-MRLC (3671; Cell Signaling Technology), α-catenin (610193; BD Transduction Labs), anti-Flag (clone M2, F1804; Sigma-Aldrich). Working dilutions were 1:50 to 1:500 for immunofluorescence and 1:500 to 1:2,000 for Western blot. One exception was the polyclonal α-catenin (α18) antibody (Yonemura et al., 2010), which was a gift from Dr. Akira Nagafuchi (Nara Medical University, Japan), used at 1:4000 for immunofluorescence.

The secondary antibodies used in the present study were HRP anti–mouse (711-035-150; Jackson ImmunoResearch Laboratories), HRP anti–rabbit (711-035-152; Jackson ImmunoResearch Laboratories), Clean-Blot HRP IP Detection Reagent (21230; Thermo Fisher Scientific), Alexa Fluor 488 anti–mouse (A21202, A11029; Invitrogen), Alexa Fluor 488 anti–rabbit (A21206, A11034; Invitrogen), Alexa Fluor 488 anti–rat (A11006; Invitrogen), Alexa Fluor 594 Phalloidin (A12381; Invitrogen), Alexa Fluor 647 anti–mouse (A21236; Invitrogen), Alexa Fluor 647 anti–rabbit (A21245; Invitrogen). Working dilutions were 1:500 for immunofluorescence and 1:2,000 for Western blot. Exceptions included: Alexa Fluor 594 Phalloidin, used at 1:100 for immunofluorescence and Clean-Blot HRP used at 1:400 when blotting for PDLIM1 in IP-eluted samples.

### Immunoprecipitation

Cells were grown on 10 cm plates until fully confluent, placed on ice, washed with ice-cold PBS, and lysed with ice-cold Triton X-100 lysis buffer (150 mM NaCl, 1 mM EDTA, 50 mM Tris, pH 7.4, and 1% Triton X-100) containing 2X protease (Protease Inhibitor Cocktail III, RPI) and phosphatase inhibitors (Halt Phosphatase Inhibitor Cocktail, Thermo Scientific). Six 10 cm plates were used per IP. Prior to cell lysis, 7 µg of antibody or IgG control (011-000-003; Jackson ImmunoResearch Laboratories) was incubated with 50 μl Protein G Dynabeads (Invitrogen) in 200 µl PBS with 0.02% Tween-20 for 10-12 hrs at 4°C with constant end-to-end rotation. For the AGO2 IP, 8 µg of antibody or IgG isotype control (ab18443; Abcam) were used. After washing the antibody-conjugated beads three times with IP lysis buffer, cell lysates were incubated with the bead-conjugated antibodies 12 hrs at 4°C with constant end-to-end rotation. Beads were then washed three times with IP lysis buffer and eluted using 50 mM DTT (Sigma-Aldrich) and 0.5% SDS in lysis buffer at 58°C for 30 min, with constant agitation. Eluted proteins were separated by SDS-PAGE and analyzed by immunoblotting as described below.

### RNA immunoprecipitation (RNA-IP) and qRT-PCR

Cells were grown on 10 cm plates until fully confluent. Cells were then immediately placed on ice, washed once with sterile ice-cold PBS, and lysed with ice-cold, filter-sterilized, IP lysis buffer (150 mM NaCl, 1 mM EDTA, 50 mM Tris pH 7.4, 1 % Triton X-100), containing 2x amount of protease (cocktail III, RPI) and phosphatase inhibitors (Pierce), as well as 100 U/ml RNAse Inhibitor (RNAsin; Promega). Cells were scraped, left for 20 min on ice and lysates were cleared up by full-speed centrifugation for 5 min at 4°C and pre-cleared with Protein G Dynabeads (Life) for 1h at 4°C with constant end-to-end rotation. Lysate from a total of 2 x 10 cm plates was used per IP. In parallel, 4 μg of rabbit AGO2 antibody (ab156870; Abcam) or rabbit IgG control (011-000-003; Jackson ImmunoResearch Laboratories) were incubated O/N with 40 μls Protein G Dynabeads (Invitrogen) in 300 μls sterile 1xPBS + 0.02 % Tween and then washed 3x with IP lysis buffer. Pre-cleared lysates were incubated with the beads-conjugated antibodies for 12 hours at 4°C with constant end-to-end rotation, with the exception of 100 μls that were kept at -20°C as the total lysate control. Beads were then washed 3x with ice-cold IP lysis buffer (sterile) and immunoprecipitated complexes were eluted using 100 μls elution buffer composed of: IP lysis buffer + 2x protease + 2x phosphatase inhibitors + 100U/ml RNAsin inhibitors + 50 mM dithiothreitol (DTT) + 0.5 % SDS (sterile) at 58°C for 30 min. Eluates were then transferred in fresh tubes and RNA extraction was performed by adding 500 μls Trizol (Invitrogen) and by using the Trizol Plus Total Transcriptome Isolation protocol of the PureLink RNA mini kit (Ambion - Life Technologies). Final RNA concentrations were determined using a BioDrop spectrophotometer. RNA was converted to cDNA using the High Capacity cDNA Reverse Transcriptase Kit (Applied Biosystems). qPCR reactions were performed using the Taqman FAST Universal PCR master mix (Applied Biosystems), in a BioRad CFX96 Touch/Connect real time quantitative PCR machine. Data were analyzed and normalized using the Sigma RIP-qRT-PCR Data Analysis Calculation Shell, associated with the Sigma Imprint RIP kit (http://www.sigmaaldrich.com/life-science/epigenetics/imprint-rna.html) and the references therein. TaqMan assays used (Applied Biosystems, cat# 4427975): hsa-miR-24, 000402; hsa-miR-200c, 002300; hsa-miR-203a, 000507.

### Immunoblotting

Whole-cell extracts were obtained using RIPA buffer (50 mM Tris, pH 7.4, 150 mM NaCl, 1% NP-40, 0.5% deoxycholic acid, and 0.1% SDS) supplemented with protease and phosphatase inhibitors. Lysates were homogenized through a 29-G needle and cleared by full-speed centrifugation for 5 min. Protein quantification was performed using a Pierce BCA Protein Assay (Thermo Fisher Scientific). Protein extracts were mixed with Laemmli sample buffer and separated by SDS-PAGE using 4-20% TGX gels (Bio-Rad) and transferred to 0.2 µm nitrocellulose membranes (Bio-Rad) with the Trans-Blot Turbo Transfer System (Bio-Rad). Membranes were blocked and blotted in 3% milk according to standard protocols. Signals were detected by luminescence using Pierce ECL (Thermo Fisher Scientific) and a ChemiDoc Imaging System (Bio-Rad).

### Immunofluorescence

Cells were grown in 12 well plates on 18 mm on sterile glass coverslips until they reached full confluence. Cells were washed once with PBS and fixed with either: a) 100% methanol (Thermo Fisher Scientific) at -20°C for 7 min; or b) 3.7% formaldehyde (Electron Microscopy Sciences) in PBS at RT for 20 min. Following 3.7% formaldehyde fixation, cells were washed 3 times with PBS + glycine (10 mM) and permeabilized for 7 min using 0.15% Triton X-100 in PBS. Cells counterstained with phalloidin to visualize F-actin were 3.7% formaldehyde fixed, while most all other cells were fixed with 100% methanol. Cells immunostained using the α18 antibody (Yonemura et al., 2010) were fixed with 1% formaldehyde in 0.1M HEPES (Gibco) at RT for 20 min, washed 3 times with PBS + glycine (10 mM), and permeabilized for 12 min using 0.15% Triton X-100 in PBS.

Cells were blocked with serum free Protein Block reagent (Dako) at RT for 1hr and stained with primary antibodies diluted in Antibody Diluent (Dako) overnight at 4°C. Cells were then washed three times with PBS, stained with the fluorescent-labeled secondary antibodies for 1 hr at RT, washed three times with PBS, co-stained with DAPI (Sigma-Aldrich), and mounted (Aqua-Poly/Mount; Polysciences).

### Calcium switch and cytoskeletal inhibitor treatments

Caco2 cells were grown on coverslips until confluence. In Latrunculin A (LatA)-treated calcium switch assays, cells were pretreated with either DMSO (Fisher Scientific) or 10 µM LatA (Thermo Fisher Scientific) for 10 min, washed three times with calcium-free PBS, and incubated in calcium-free Caco2 medium (S-MEM; Gibco), supplemented with glutamine, 10% FBS, sodium pyruvate, NEAA, and containing 4 mM EGTA for 30 min, until cells were rounded. Calcium-free EGTA-containing medium also contained either DMSO or 10 µM LatA. Cells were then washed three times with PBS, returned to regular Caco2 medium, again with either DMSO or LatA, and fixed for immunofluorescence at the indicated time points. In Blebbistatin or Y-27632-treated calcium switch assays, there was no pre-treatment prior to incubation with calcium-free EGTA-containing Caco2 medium. Treatment was introduced in recovery, following cell rounding. In recovery, regular Caco2 medium contained either 50 µM (±)-Blebbistatin (Calbiochem)/DMSO control or 30 µM Y-27632 (Calbiochem)/water control. For static cytoskeletal inhibitor treatments, confluent cells were treated with 50 µM Blebbistatin for 3 hrs or 20 µM Y-27632 for 2 hrs.

### Proximity Ligation Assay (PLA)

Confluent cells were fixed by using 4% paraformaldehyde (Electron Microscopy Sciences) and permeabilized with 0.2% Triton-X (Thermo Fisher Scientific) in PBS at room temperature. Coverslips were blocked using a blocking reagent (Dako) for 1 h at room temperature and probed with primary antibodies diluted in antibody diluent (Dako) overnight at 4°C. Primary antibodies used were anti-Flag (clone M2; F1804; Sigma-Aldrich) and Myosin IIB (909901; Biolegend) at 1:500 dilution. Coverslips were then washed with Duolink® In Situ Wash Buffer-A (Sigma-Aldrich; DUO82049) and incubated with a Duolink® Anti-mouse plus PLA probe (Sigma-Aldrich) together with a Duolink® Anti-rabbit minus PLA probe (Sigma-Aldrich) at 37°C, for one hour, in a humidified chamber. Coverslips were then washed with Duolink® In Situ Wash Buffer-A (Sigma-Aldrich), incubated with ligation solution (Sigma-Aldrich) for 30 min at 37°C, washed with Duolink® In Situ Wash Buffer-A, and incubated with polymerase solution (Sigma-Aldrich) for 100 min at 37°C. Coverslips were then washed with Duolink® In Situ Wash Buffer-B (Sigma-Aldrich), then with Duolink® In Situ Wash Buffer-A and counterstained with Alexa flour 488 conjugated E-cadherin antibody (3199S; Cell Signaling Technology) for 1 h at room temperature. Coverslips were eventually washed with Duolink® In Situ Wash Buffer-B and with Duolink® In Situ Wash Buffer-A before being mounted with Duolink® In Situ Mounting Medium with DAPI (Sigma-Aldrich). Coverslips were imaged using a Leica SP8 confocal microscope under 63x objective with an additional 1.5x zoom, with Z-stack acquisition intervals at 0.5–1 μm, image resolution at 1024 x 1024 pixels, 600 Hz scan speed, and scan line average at 4.

### Gene ontology enrichment analysis

The full published dataset of the 607 PLEKHA7-interacting proteins (publicly available at http://www.imexconsortium.org/; identifier: IM-25739) (Kourtidis et al., 2017b) was analyzed using the Gene Ontology (GO) Enrichment analysis online tool (http://geneontology.org) (Ashburner et al., 2000; Gene Ontology, 2021; Mi et al., 2019). Annotation version and release date of the PANTHER database used was: Reactome version 65, released 2020-11-17. Statistical analysis was performed using the PANTHER Overrepresentation Test tool. Test type used was Fisher’s exact test with false discovery rate (FDR) correction. Annotation datasets used for the analysis were the PANTHER GO-slim Molecular Function and the PANTHER GO-slim Biological Process.

### Imaging, image processing, analysis, and statistics

Images were acquired using Leica SP5 and SP8 confocal microscopes with 63x Plan-Apochromat 1.4NA DIC oil immersion objectives (Leica) and 405 nm, 488 nm, 594 nm, and 633 nm lasers. Image acquisition was done using Leica Application Suite Advanced Fluorescence X software at 1024 x 1024 resolution and with 0.4 µm intervals along the z*-*axis. For super resolution imaging, Z-series stacks were acquired in 0.4 µm intervals on a Zeiss LSM 880 NLO Quasar point scanning confocal microscope using the Airyscan detector, a 63x Plan-Apochromat 1.4NA DIC oil immersion objective (Zeiss) and the 405 nm, 488 nm, 561 nm, and 633 nm laser lines at 2024 x 2024 resolution, with 1.8X zoom. The Zeiss Zen 2.3 (black edition) software was used to control the microscope, adjust spectral detection, and for processing of the Airyscan raw images.

To allow for comparison, the same imaging parameters were used across conditions for each set of experiments. Unless otherwise noted, images shown are single Z-slices. For quantifications of confocal and super resolution images, maximum intensity projections of the two most apical Z-slices were used to account for uneven cell thicknesses and ensure full representation of all apical cell-cell junctions throughout each field. Quantifications were done using Fiji (Schindelin et al., 2012) (National Institutes of Health). Individual borders for measuring signal intensity or junction straightness were manually and randomly selected based on apical signals, using apical E-cadherin as a reference in most cases. Integrated fluorescence intensity values were measured using Fiji from drawing either: a) 6 µm lines tracing cell-cell borders, when assessing junctional marker fluorescence intensity, normalized to fluorescence intensity averages of equal number of 6 µm lines drawn randomly throughout the cytoplasm, to account for staining fluctuations and/or total protein levels; b) 6 or 3 µm (e.g. for the super-resolution images) lines perpendicular to cell-cell borders, when assessing both fluorescence intensity, as well as distribution of this marker across cell-cell junctions; c) 8 × 3 µm boxes centered on cell-cell junctions when using the α18 antibody, to include jagged junctions. To measure junctional straightness, the length of a line tracing the cell border from one tricellular junction to the next was compared with the length of a straight line drawn directly from one tricellular junction to another.

For all measurements, sample size and related statistics are indicated in the respected figure legends. In general, we used: a) unpaired two-way *t* tests when comparing two groups; b) one-way ANOVA when comparing three or more groups; and c) two-way ANOVA when more than one parameters were considered (e.g. fluorescence intensity and distance). Statistics and graphs were all performed using Prism 10 (GraphPad). All experiments, including gene knockdowns and immunoprecipitation experiments, were performed in at least three independent experiments, unless otherwise noted, and representative images are shown.

## ACKNOWLEDGEMENTS

This work was supported by NIH grants R01 DK124553, R21 CA246233, P20 GM130457, 3P20 GM130457-04S1, P30 DK123704, P20 GM103499-21S1, UL1 TR001450 (MUSC South Carolina Clinical & Translational Research Institute Discovery Grant), and P30 CA138313 (Hollings Cancer Center Pre-Clinical & Clinical Concepts Pilot Award) to AK. MCB was supported by NIH training grants TL1 TR001451 and UL1 TR001450. AR was supported by NIH training grants TL1 TR001451, UL1 TR001450 and T32 5T32DK124191. AD was supported by the Abney Graduate Fellowship Award, Hollings Cancer Center (P30 CA138313), MUSC. CK was supported by NIH training grant 5T32 DE017551.

We would like to thank Dr. Akira Nagafuchi, Nara Medical University, Japan, for kindly providing the α18 antibody; Drs. Christiana Kappler and Stephen Duncan, Cell Models Core, MUSC Center of Biomedical Research Excellence (COBRE) in Digestive & Liver Disease (CDLD; NIH P20 GM130457) and MUSC Digestive Disease Research Center (DDRC; NIH P30 DK123704) for support with CRISPR/Cas9 reagents; Drs. Li Li and John Lemasters, Advanced Imaging Core, CDLD and DDRC (NIH P20 GM130457 and P30 DK123704) for support with super resolution microscopy; Dr. David Turner and the shRNA Technology Shared Resource, Hollings Cancer Center, MUSC (NIH P30 CA138313) for providing shRNA constructs.

The authors declare no competing financial interests.

Author contributions: Conceptualization: MCB, AK; data acquisition: MCB, JNM, AR, DWJ, ACD, CK, MED; data analysis: MCB, JNM, AR, DWJ, ACD, AK; writing - original manuscript: MCB, AK; writing - review, editing, revised manuscript: AK

---

**FIGURE S1.**
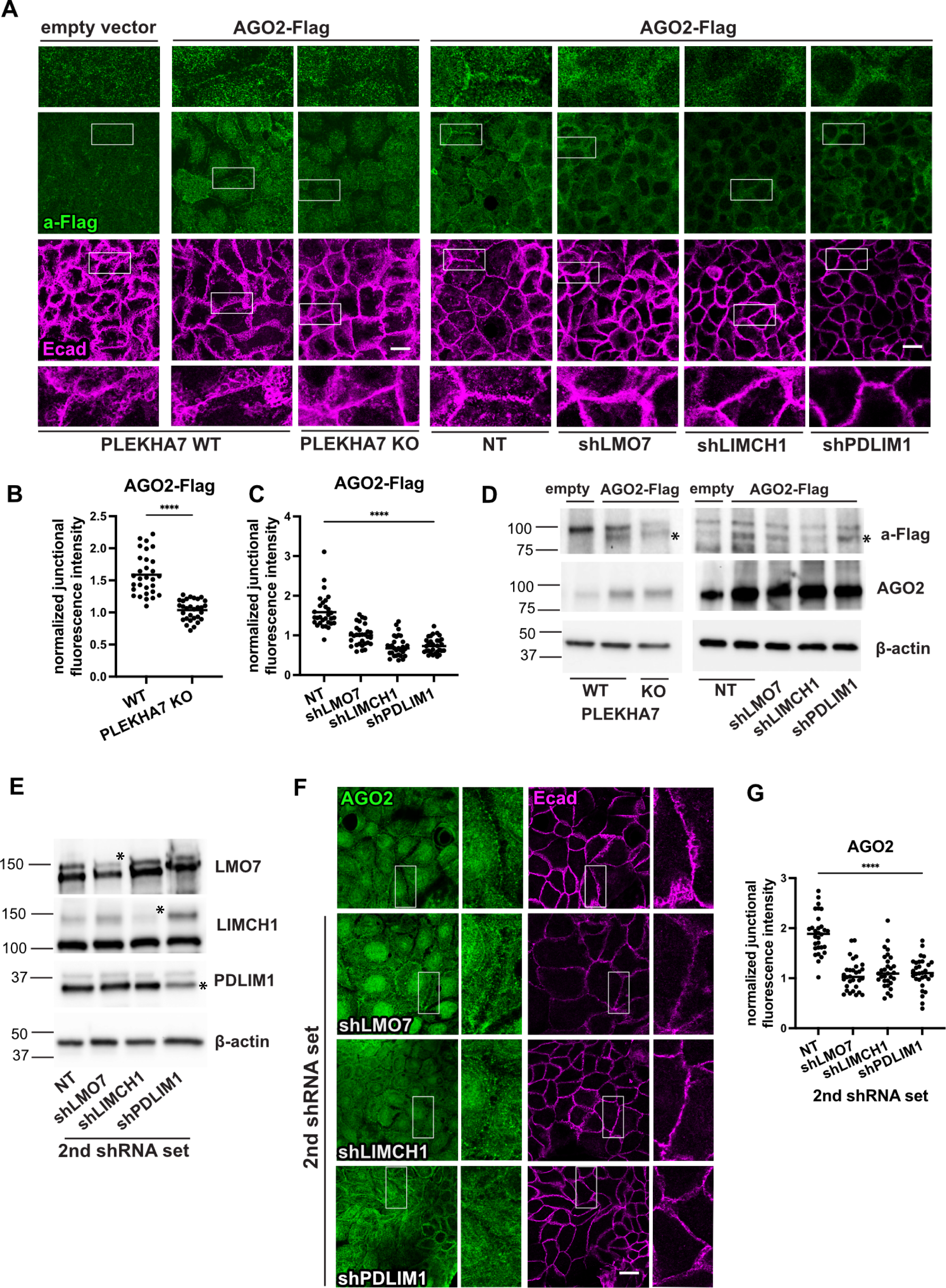
Regulation of junctional localization of AGO2 demonstrated by an ectopically expressed AGO2-Flag construct and a second set of shRNAs targeting LMO7, LIMCH1, and PDLIM1. (A) Control (WT) or PLEKHA7 knockout (KO) and non-target (NT) shRNA control or LMO7, LIMCH1, and PDLIM1 shRNA-mediated knockdown (shLMO7, shLIMCH1, shPDLIM1, respectively) Caco2 cells were stably transduced with an empty vector control or an AGO2-Flag – expressing construct and subjected to immunofluorescence using an anti-Flag antibody and E-cadherin (Ecad). (B-C) AGO2-Flag junctional fluorescence intensity from (A) was normalized to cytoplasmic and quantified from n=30 cell-cell junctions (10 junctions/field) representative of three independent experiments; statistical analyses were performed using unpaired two-way *t* test (B) or one-way ANOVA test (C); ****, P < 0.0001. (D) Immunoblotting of (WT) or PLEKHA7 knockout (KO) and of non-target (NT) shRNA control or LMO7, LIMCH1, and PDLIM1 shRNA-mediated knockdown (shLMO7, shLIMCH1, shPDLIM1, respectively) Caco2 cells transduced with empty vector or the AGO2-Flag construct, for a-Flag, AGO2; β-actin is the loading control. Molecular masses (kD) are indicated on the left. (E) Immunoblotting of control (NT) or LMO7, LIMCH1, and PDLIM1 knockdown Caco2 cells using a second set of shRNAs for each target (shLMO7, shLIMCH1, shPDLIM1 – 2^nd^ shRNA set); asterisks indicate the specific LMO7, LIMCH1, PDLIM1 bands lost by shRNA targeting. β-actin is the loading control; molecular masses (kD) are indicated on the left. (F) Immunofluorescence of NT, shLMO7, shLIMCH1, shPDLIM1 Caco2 cells for AGO2 and E-cadherin (Ecad). Insets in all cases are marked by white rectangles and are 3X magnification of the original image. (G) AGO2 and PLEKHA7 junctional fluorescence intensity from (F) was normalized to cytoplasmic and quantified from n=30 cell-cell junctions representative of three independent experiments; statistical analyses were performed using one-way ANOVA tests; ****, P < 0.0001. All scale bars = 20 µm.

**FIGURE S2.**
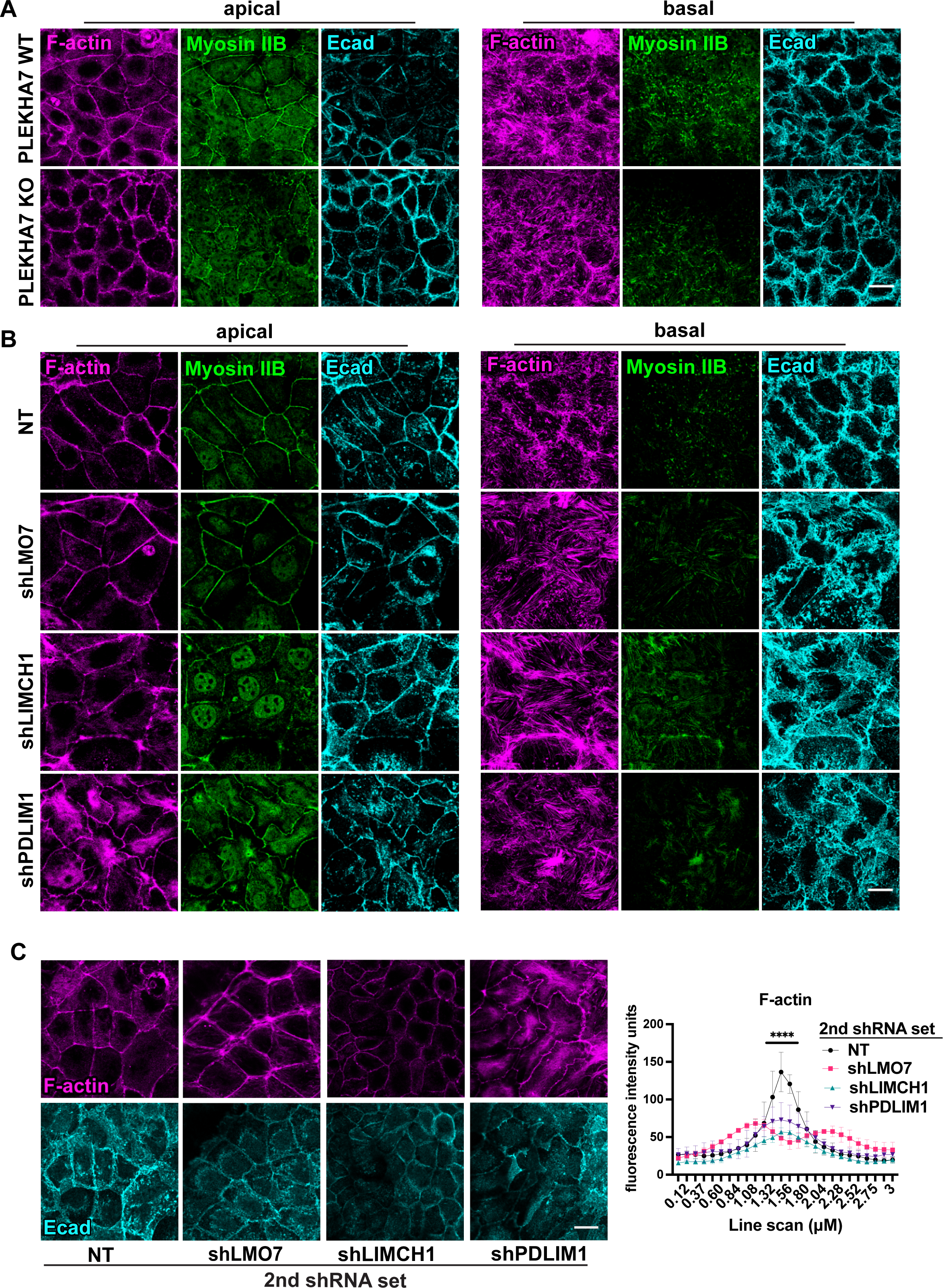
Actomyosin organization is disrupted in PLEKHA7 knockout and LMO7, LIMCH1, PDLIM1 knockdown cells, also by using different shRNA sets. Immunofluorescence and confocal imaging of (A) wild type (WT) or PLEKHA7 knockout (KO) Caco2 cells, and (B) of control (NT) or LMO7, LIMCH1, and PDLIM1 knockdown (shLMO7, shLIMCH1, shPDLIM1) Caco2 cells, accompanying super resolution imaging (see Fig. 4). Representative apical and basal Z-slices are shown. (C) Immunofluorescence and confocal imaging of control (NT) or LMO7, LIMCH1, and PDLIM1 knockdown Caco2 cells using a second shRNA for each target (shLMO7, shLIMCH1, shPDLIM1 – 2^nd^ shRNA set) for F-actin and E-cadherin (Ecad). F-actin fluorescence intensity was measured using 3 µm line scans drawn perpendicular to cell-cell junctions from n=30 cell-cell junctions (10 junctions/field) representative of three independent experiments; statistical analyses were performed using two-way ANOVA; error bars represent mean ± SD; ****, P < 0.0001. Scale bars = 20 µm.

**FIGURE S3.**
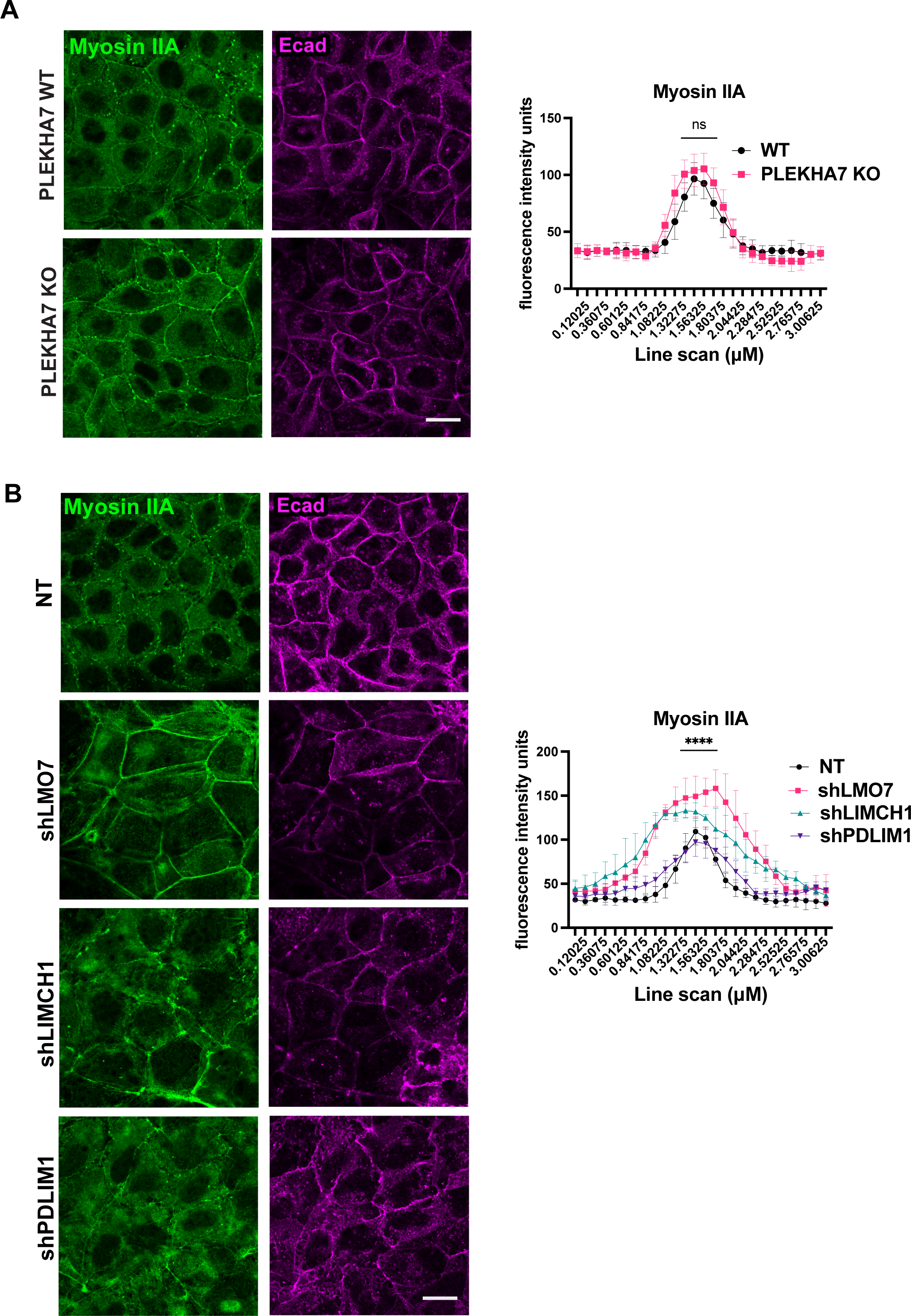
PLEKHA7 knockout and LMO7, LIMCH1, PDLIM1 knockdown do not have uniform effects on Myosin IIA distribution. Immunofluorescence and confocal imaging of (A) wild type (WT) or PLEKHA7 knockout (KO) and (B) of control (NT) or LMO7, LIMCH1, and PDLIM1 knockdown (shLMO7, shLIMCH1, shPDLIM1) Caco2 cells, for Myosin IIA and E-cadherin (Ecad). Myosin IIA fluorescence intensity was measured using 3 µm line scans drawn perpendicular to cell-cell junctions from n=30 cell-cell junctions representative of two independent experiments; statistical analyses were performed using two-way ANOVA; error bars represent mean ± SD; ****, P < 0.0001; ns, non-significant. Scale bars = 20 µm.

**FIGURE S4.**
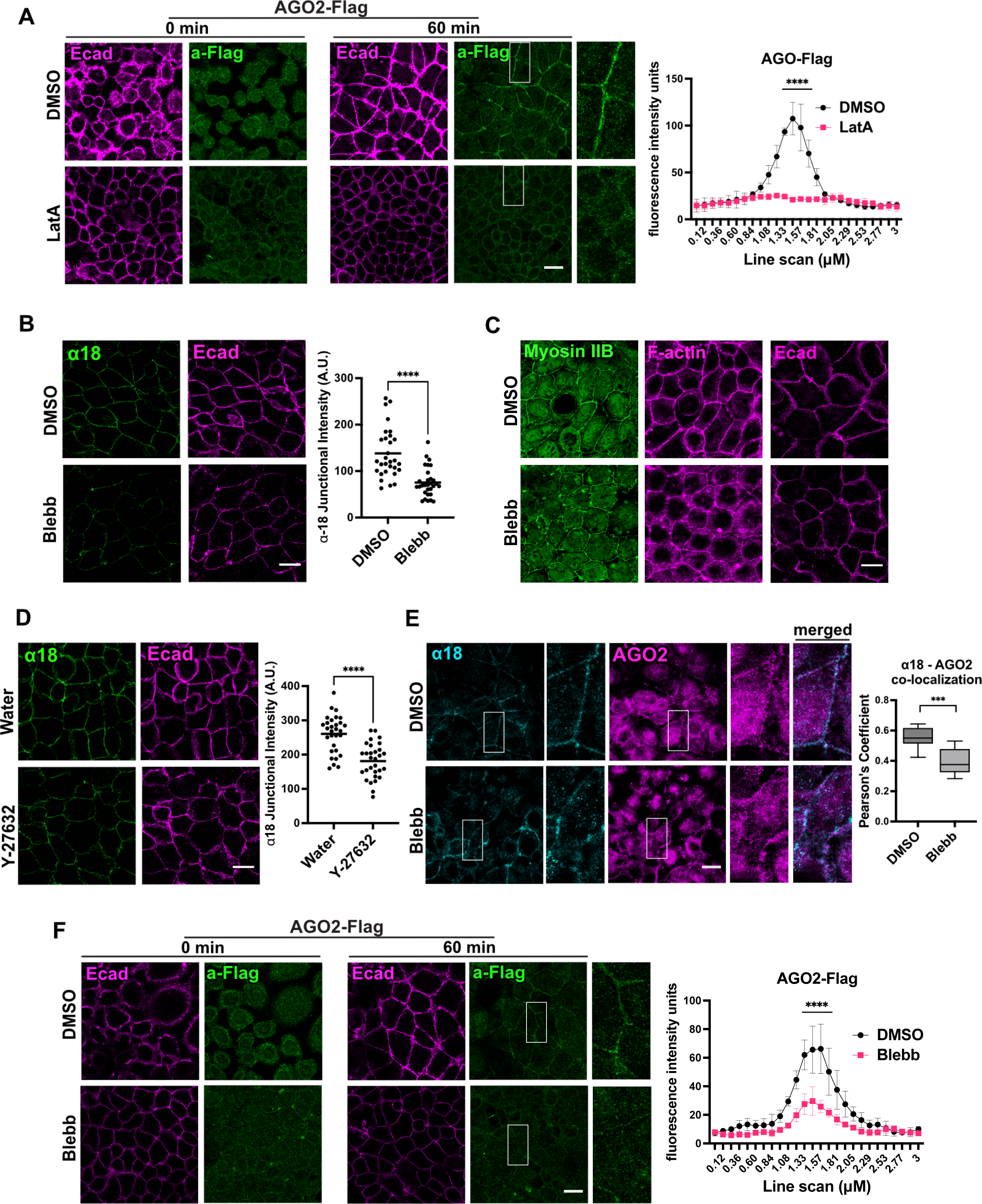
Disruption of actin polymerization and tension result in loss of junctional recruitment of both the endogenous and of ectopically expressed AGO2. (A) Immunofluorescence of Caco2 cells stably transduced with an AGO2-Flag construct for a-Flag and E-cadherin, fixed at indicated time points throughout a calcium switch assay. Images at 0 min indicate Ca^2+^ depleted cells immediately before Ca^2+^ reintroduction and at 60 min time point are following Ca^2+^ re-addition. Cells were pre-treated and maintained in either DMSO vehicle (control) or 10 µM Latrunculin A (LatA) throughout the calcium switch assay. Insets are marked by white rectangles and are 3X magnification of the original image. Fluorescence intensity of 3 µm line scans drawn perpendicular to cell-cell junctions was measured for a-Flag at the 60 min time point upon recovery from n=30 cell-cell junctions (10 junctions/field) representative of three independent experiments; statistical analysis was performed using two-way ANOVA; error bars represent mean ± SD; ****, P < 0.0001. (B-D) Immunofluorescence of the α18 α-catenin antibody, or of F-actin and Myosin IIB in Caco2 cells, treated with DMSO vehicle or 50 µM Blebbistatin (Blebb) for 3 h (B-C), or with water vehicle or 20 µM of the ROCK inhibitor Y-27632 for 2 h (D). For fluorescence intensity quantifications, n= 30 individual junctions/treatment group obtained by measuring 10 cell-cell junctions/field from 3 fields. Statistical analysis was performed using an unpaired two-way *t* test; ****, P < 0.0001. (E) Immunofluorescence of AGO2 and α18 α-catenin at 60 min upon recovery in a calcium switch assay, during which DMSO or Blebbistatin were included in the calcium-containing recovery medium. AGO2 and α18 co-localization was measured using Pearson’s Coefficient from n=10 cell-cell junctions (5 junctions/field) representative of two independent experiments. Error bars represent mean ± SD; ***, P < 0.001. (F) Immunofluorescence of Caco2 cells stably transduced with an AGO2-Flag construct for a-Flag and E-cadherin, during a calcium switch assay in which DMSO or Blebbistatin were included in the calcium-containing recovery medium, fixed at indicated time points. Images at 0 min indicate Ca^2+^ depleted cells immediately before Ca^2+^ reintroduction and at 60 min time point are following Ca^2+^ re-addition. Insets are marked by white rectangles and are 3X magnification of the original image. Fluorescence intensity of 3 µm line scans drawn perpendicular to cell-cell junctions was measured for a-Flag at the 60 min time point upon recovery from n=30 cell-cell junctions (10 junctions/field) representative of three independent experiments; statistical analysis was performed using two-way ANOVA; error bars represent mean ± SD; ****, P < 0.0001. All scale bars= 20 µm.

**FIGURE S5.**
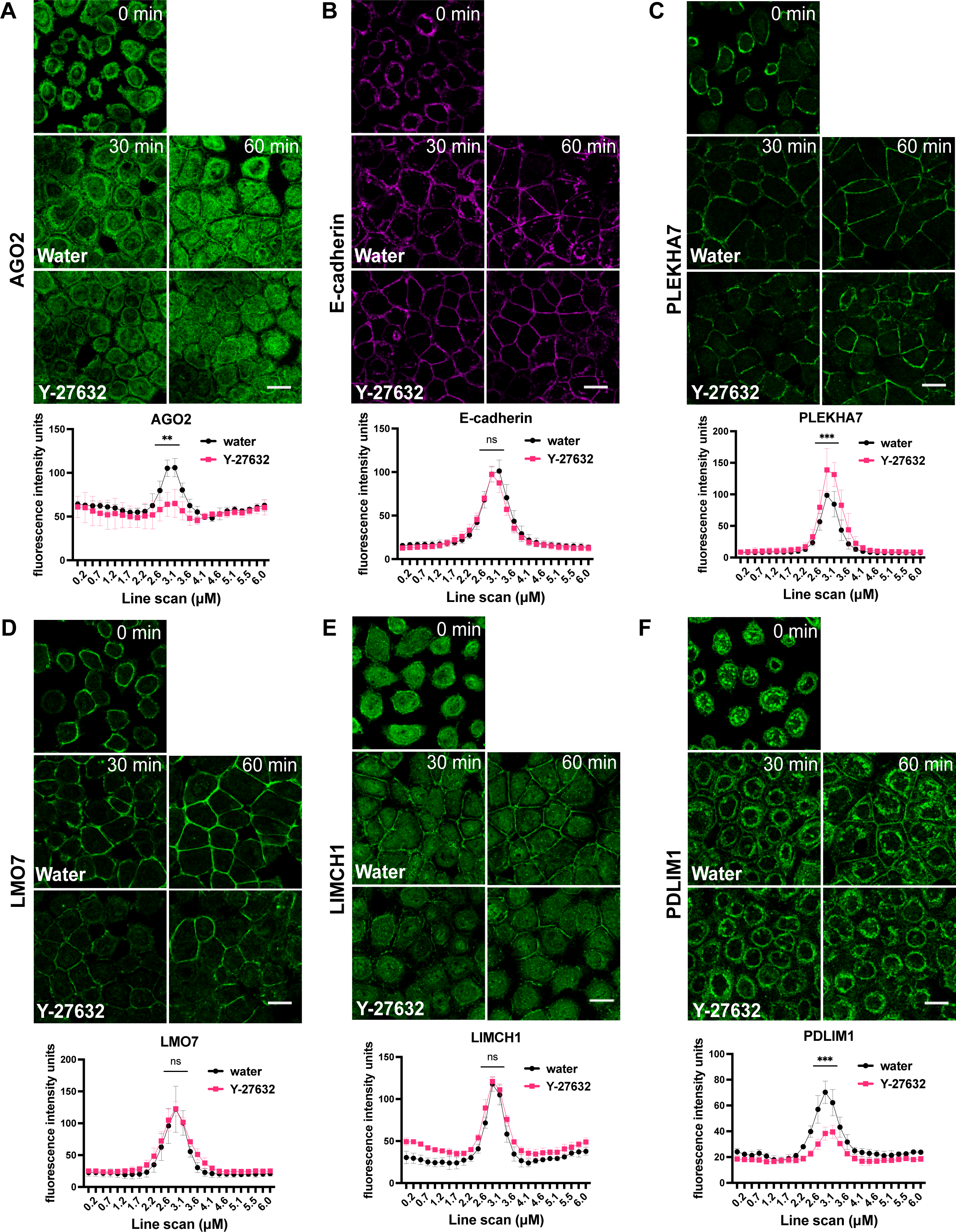
Y-27632 treatment during recovery following calcium depletion results in loss of junctional AGO2. (A-F) Immunofluorescence of a calcium switch assay in which water vehicle or 30 µM Y-27632 were included in the calcium-containing recovery medium. Cells fixed immediately before Ca^2+^ reintroduction (0 min) or 30, 60 min post Ca^2+^ addition were stained for AGO2 (A), E-cadherin (B), PLEKHA7 (C), LMO7 (D), LIMCH1 (E), and PDLIM1 (F). Fluorescence intensity of 6 µm line scans drawn perpendicular to cell-cell junctions was measured for each marker at the 60 min time point upon recovery from n=30 cell-cell junctions (10 junctions/field) representative of three independent experiments; statistical analyses were performed using two-way ANOVA tests. Error bars represent mean ± SD. ***, P < 0.001; **, P < 0.01; ns, non-significant. Scale bars = 20 µm.

**FIGURE S6.**
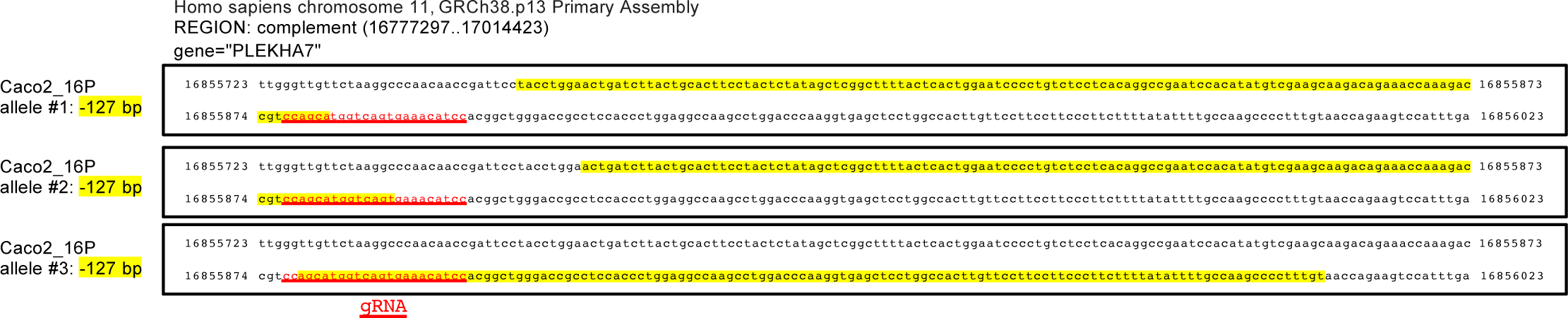
PLEKHA7 CRISPR/Cas9 knockout validation. Sequencing of the genomic region of PLEKHA7 that was targeted by CRISPR/Cas9 in the PLEKHA7 knockout Caco2 cells (clone 16P) used in the present study. The guide RNA (gRNA) sequence used is underlined. The sequenced regions were aligned against the human genome GRCh38.p13 Primary Assembly. The deleted regions detected in PLEKHA7 alleles are highlighted (Caco2 cells exhibit polyploidy).

## Notes

### Competing Interest Statement

The authors have declared no competing interest.

### Summary of Updates

Revisions include: a) revised Introduction and Results section; b) new Discussion section; c) additional data and Figures (2E; 6; 8; S1; S2C; S3; S4); d) quantifications of all immunofluorescence experiments; e) new experimentation, including constructs (AGO2-Flag), additional shRNAs, new markers examined (Myosin IIA, pMRLC), and new assays (RNA immunoprecipitation and proximity ligation)

